# Morphological signatures of neurogenesis and neuronal migration in hypothalamic vasopressinergic magnocellular nuclei of the adult rat

**DOI:** 10.1101/2023.12.10.570983

**Authors:** Limei Zhang, Mario A. Zetter, Vito S. Hernández, Oscar R. Hernández-Pérez, Fernando Jáuregui-Huerta, Quirin Krabichler, Valery Grinevich

## Abstract

Arginine vasopressin (AVP)-magnocellular neurosecretory system (AVPMNS) in the hypothalamus plays a critical role in homeostatic regulation as well as in allostatic motivational behaviors. However, it remains unclear whether adult neurogenesis exists in the AVPMNS. By using immunoreaction against AVP, neurophysin II, glial fibrillar acidic protein (GFAP), cell division marker (Ki67), migrating neuroblast markers (doublecortin, DCX), microglial marker Iba1, and 5′-bromo-2′-deoxyuridine (BrdU), we report morphological evidence that low rate neurogenesis and migration occur in adult AVPMNS in rat hypothalamus. Tangential AVP/GFAP migration routes and AVP/DCX neuronal chains as well as ascending AVP axonal scaffolds were observed. Chronic water deprivation significantly increased the BrdU+ nuclei within both the SON and PVN. These findings raise new questions about AVPMNS’s potential hormonal role for brain physiological adaptation across the lifespan, with possible involvement in coping with homeostatic adversities.

## Introduction

Adult neurogenesis is defined as the ability of specialized cells in the postnatal brain to produce new functional neurons. This phenomenon was first formally reported six decades ago using intraperitoneal injection of thymidine-H^3^ in rats and cats, which shed some light on the possibility that new neurons may be formed in forebrain structures in both rodents and carnivores, which was not considered possible previously (Altman 1962, Altman 1963, Altman and Das 1966). Today, it is well established that the subventricular zone of the lateral ventricles and the subgranular zone of the hippocampus are prominent regions of neurogenesis that occurs through the lifespan (Doetsch and Alvarez-Buylla 1996, Lois, Garcia-Verdugo and Alvarez-Buylla 1996, Zhao, Deng and Gage 2008, Ming and Song 2011, Zhou, Su et al. 2022). However, the phenomenon has been described in a large number of additional brain regions (Jurkowski, Bettio et al. 2020). For example, adult neurogenesis has also been observed in the striatum of human (Ernst, Alkass et al. 2014), in hippocampal fissure region in rats under pilocarpine insult (Zhang, Hernandez et al. 2014), and in caudate nucleus and nucleus accumbens in the rat (Das and Altman 1970). Furthermore, it has been reported in hypothalamus of several rodent and primate species (Kokoeva, Yin and Flier 2005, Kokoeva, Yin and Flier 2007, Migaud, Batailler et al. 2010, Cheng 2013, Yoo and Blackshaw 2018, Sharif, Fitzsimons and Lucassen 2021, Yao, Taub et al. 2021), suggesting that adult hypothalamic neurogenesis may serve as a mechanism for brain physiological adaptation across the lifespan, with possible roles in coping with homeostatic adversities. However, the actual sites and organization of distinct neurogenic niches in the hypothalamus remain elusive.

The hypothalamus hosts unique magnocellular neurosecretory neurons (MCN) which synthesize the neurohormone arginine vasopressin (AVP) and its homologue oxytocin (OT) (Bargmann and Scharrer 1951, Hatton 1990). Three loci of AVP-MCNs have been described, i.e., the supraoptic (SON), paraventricular (PVN), and accessory (AN) nuclei (Ugrumov 2002). Most, if not all, of the AVP-MCN located within the SON, PVN and AN emit main axons carrying AVP to the pituitary gland, via the hypothalamic-neurohypophysial tract (Falke 1991), which exerts control over body water and electrolyte homeostasis, blood pressure, and hepatic glucose metabolism (Sladek and Armstrong 1987, Cunningham and Sawchenko 1991, Ugrumov 2002, Armstrong 2004). The role of AVP-MCN in hormonal regulation of peripheral functions has been gradually established over the past century. However, a novel understanding of AVP-MCN involvement in higher brain functions, such as memory processes, emotion and motivation, complementary to their peripheral physiological role, has emerged in recent years (for reviews see (Zhang and Eiden 2019, Zhang, Hernández et al. 2021)).

In the early 1990s, Swaab and colleagues reported that the hypothalamic AVP and OT nuclei change in size and cell number at various postnatal developmental stages in the pig (van Eerdenburg, Poot et al. 1990, van Eerdenburg and Swaab 1991, van Eerdenburg and Swaab 1994). These changes were also observed upon alteration of gonadal- reproductive status (van Eerdenburg, Lugard-Kok et al. 1991). The above observations were further confirmed by a Canadian group (Rankin, Partlow et al. 2003, Raymond, Kucherepa et al. 2006), who, by using double-immunohistochemistry against proliferating-cell nuclear antigen (PCNA, a key factor in DNA replication) and either AVP or OT, found evidence suggesting that AVP or OT neurons could undergo postnatal cell division according to physiological demands, such as the commencement of sexual maturity and lactation. While these observations went largely overlooked, it was independently reported in 2004 that progenitor cells could be isolated from adult rat hypothalamus, which readily differentiated into mature neurons expressing various neuropeptides, among them AVP and OT (Markakis, Palmer et al. 2004). However, little is still known about the phenomenon of AVP or OT neurons generated by neurogenesis in the adult brain with respect to both, neuroanatomical organization and potential functions.

To test the hypothesis that AVP system could undergo neurogenesis and neuronal dispersion from adult rat hypothalamic magnocellular loci, we conducted a systematic histological examination of serial rat brain sections in *coronal, sagittal, horizontal and septo- temporal-oblique* orientations from male and female, juvenile to old (18 months) rats. Here, we report evidence, which strongly suggests that some AVP neurons are generated by adult neurogenesis and migrate following blood vessels, glial processes and axons of other AVP neurons. We challenged rats with chronic water deprivation every other day, and administered for two weeks intraperitoneal 5-bromo-2’-deoxyuridine (BrdU), a synthetic nucleoside analogue with a chemical structure similar to thymidine, commonly used to assess cell proliferation in living tissues (Altman 1962, Altman and Das 1966, Lieber, Shaw and Van Waes 1978). Immunohistochemical analysis revealed BrdU-immunoreactive (ir) large round-shaped nuclei inside the AVP-magnocellular loci, with some of these paired as twin-BrdU-ir nuclei, indicating ongoing neurogenesis in these areas. We also assessed the co-expression of AVP and the migrating neuroblast marker doublecortin (DCX) and found numerous DCX neurons within the SON and PVN with some of these also expressing AVP. Our data suggest that AVP neurons in rat hypothalamic SON and PVN may undergo low-rate neurogenesis during adult life and that some of them may migrate to adjacent subcortical regions, as evidenced by the presence of AVP cell chains, which might be guided by axon-scaffolds.

## Results and specific discussion

### AVP-ir SON and PVN dispersed to other subcortical regions: DCX immunopositive migrating cell chains guided by axon scaffolds

In order to describe the morphology AVP-ir in the rat brain, we analyzed serial sections cut in coronal, sagittal, horizontal, and septo- temporal (oblique) orientations. The immunohistochemical reactions were performed in every third sequential section and we used a combination of two rabbit anti-vasopressin antibodies (gift from Prof. Ruud Buijs (Buijs, Pool et al. 1989) and Peninsula laboratories international, T4563, see Tab. 1). The rationale for the use of a combination of two anti-vasopressin antibodies was to enhance phenotypic characterization by optimizing labeling of the AVP system across changes in the molecular form of vasopressin as prohormone processing ensues.

**Table 1.** Primary antibodies used and their dilutions.

| Antibody to | Host | Dilution | Source |
| --- | --- | --- | --- |
| [Arg8]-vasopressin | rabbit | 1:2000 | Peninsula laboratories international, T4563 |
| [Arg8]-vasopressin | rabbit | 1:2000 | Prof. R. M. Buijs, UNAM, Mexico |
| Neurophysin II | mouse | 1:250 | Prof. Harold Gainer, NINDS/NIH, Bethesda |
| Bromo-2-deoxyuridine (BrdU) | rat | 1:100 | Accurate Scientific, NY OBT0030 |
| Ki67 | rabbit | 1:250 | Biocare Medical, SP6 |
| Glial fibrillary acidic protein (GFAP) | rabbit | 1:1000 | Biocare Medical, 901-040-031511 |
| Neuronal nuclear antigen (NeuN) | mouse | 1:2000 | Millipore, MAB377 |
| Doublecortin (DCX) | goat | 1:1000 | Santa Cruz, SC-8066 |
| Iba1 | rabbit | 1:500 | Biocare, CP 290A |

Selective planes which could best illustrate the phenomenon of adult neuronal tangential migrations occurring in AVP-ir populations are presented in figures 1 to 5 and supplemental figure 1 with detailed description in the figure legends. Sex and age of animals from which the series were obtained are also indicated in the legends.

**Fig. 1.**
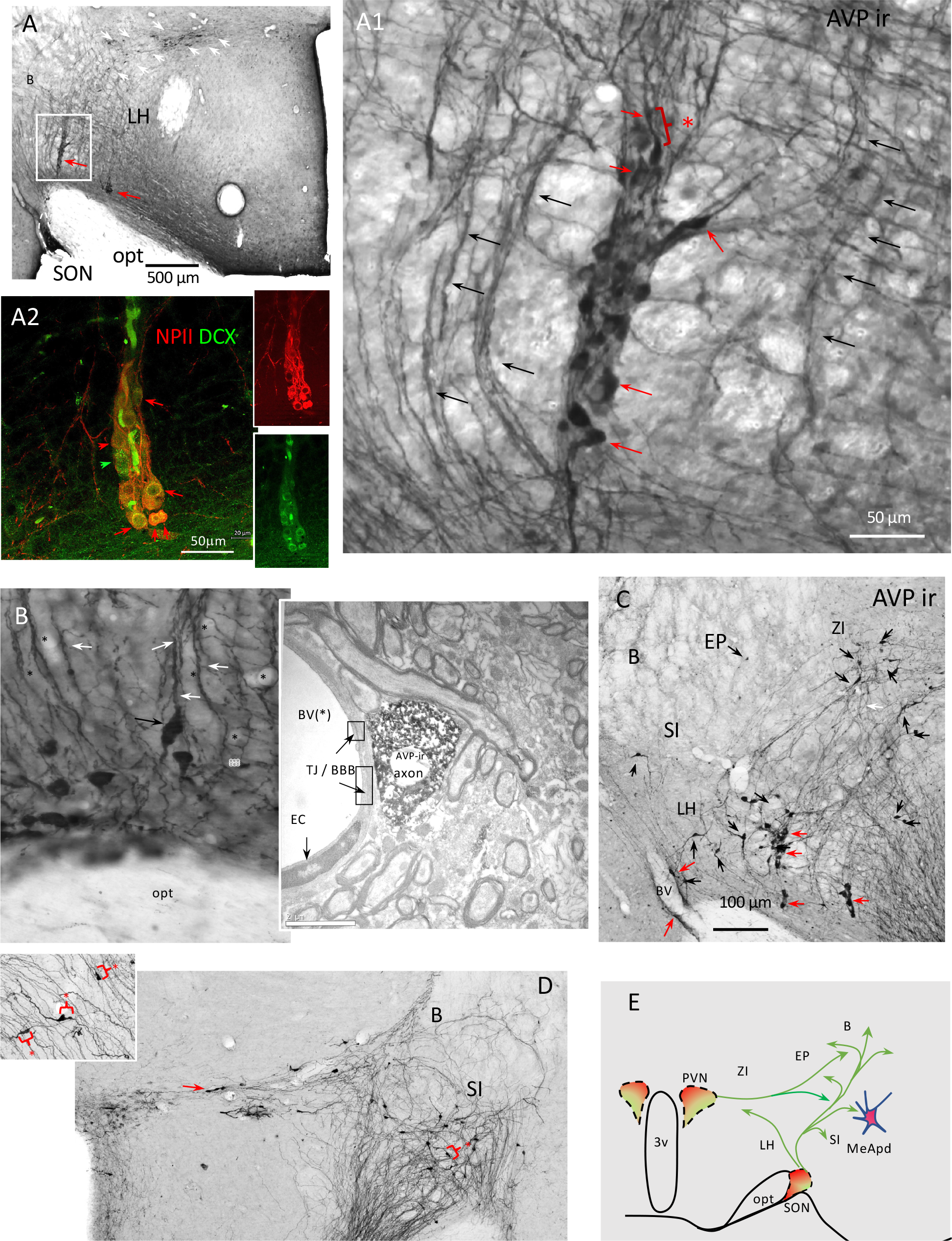
**Network of AVP-ir cell chains and axon scaffolds in the adult rat hypothalamus (1/5) emerged from supraoptic nucleus (SON) - coronal view through serial sections from an adult (4 month old) male Wistar rat. (**A) and (A1): AVP-ir cell-chains (red arrows) and axonal scaffolds (black arrows) emerged from SON^AVP^. AVP-ir cells (red arrows) showed tangential migration morphology that is mainly bipolar, with leading process extension and swelling formation for eventual somal translocation (Kaneko, Sawada and Sawamoto 2017) (white asterisk and bracket show a cell with somal translocation, as an example). The cell- chains have been previously characterized as neuronal precursors in migration (Doetsch and Alvarez-Buylla 1996). The cell-chain (A2) showed co-expression of neuroblast marker doublecortin (DCX, green) and neurophysin II (NPII, red), a carrier protein for vasopressin. Panel (B) shows the AVP-ir axonal scaffolds (white arrows). An AVP-ir cell with axon swelling and somal translocation phenotype (black arrow) seemed to “climb” into the AVP-ir scaffolds. The proximity of AVP-ir axon with blood vessels (indicated by black *) was frequently observed. A transmission electron microscopical picture with AVP IHC reaction taken from preoptic area of the rat hypothalamus is shown as (B) inset. Tight junctions (TJ) of the brain-blood-barrier (BBB) formed by processes of endothelium cells (EC) are indicated. The AVP-ir axons are separated from the lumen of blood vessels at ultra- structural level. C: presence of AVP-ir cells with tangential migration morphology in entopeduncular nucleus (EP) and lateral hypothalamus (LH). D: AVP-ir cell chain leaving the PVN (red arrow) with lateral-tangential movement morphology. Substantia innominata (SI), nucleus basalis of Meynert (B) are innervated with AVP axon scaffolds and disperse AVP-ir cells. Inset shows bipolar neurons with swelling formation for eventual somal translocation, indicated with red asterisk and brackets. E: charting symbolizing the observed migration routes from coronal samples (green arrows). Abbreviations, B: nucleus basalis of Meynert, ZI: zona incerta, 3V: third ventricle.

Figure 1 shows examples from rat brain coronal section series with an emphasis on the supraoptic nucleus (SON). AVP-ir cell-chains emerging from the SON can be clearly seen (Fig. 1, panels A & A1). Their peculiar formation and morphology first led us to hypothesize that they are putative neuronal precursors (indicated with red arrows). Indeed, cells in chains are classical identifying features for neuronal precursors in migration (Doetsch and Alvarez-Buylla 1996). The AVP-ir cells showed tangential migration morphology that is mainly bipolar, with leading process extension and swelling formation for eventual somal translocation (Kaneko, Sawada and Sawamoto 2017) (for instance, the cell indicated with red asterisk and bracket in A1). Importantly, these cell-chains (A2) showed co-expression of neurophysin II (NPII, the carrier protein for AVP; in red) and the neuroblast marker doublecortin (DCX, green). Fig. 1 panel (B) shows the AVP-ir axonal scaffolds with AVP-ir cells putatively climbing along them. The AVP-ir axons in this region (preoptic area of the rat hypothalamus) are often found in close proximity with blood vessels, as shown in the electron photomicrograph Fig. 1, B-inset. Tight junctions (TJ) formed by processes of endothelium cells can be clearly seen, which indicates that the AVP-ir axons are separated from the lumen of blood vessels by cells of the blood-brain-barrier. AVP-ir neurons with tangential migrating phenotype were further observed in entopeduncular nucleus (EP; Fig. 1 C, red arrows), again putatively climbing along AVP-ir axon scaffolds (Fig. 1 C, black arrows). Panel D shows AVP-ir cell chain leaving the PVN (red arrow) with lateral-tangential movement morphology. Substantia innominata (SI), nucleus basalis of Meynert (B) are innervated with AVP axon scaffolds and disperse AVP-ir cells. Inset shows bipolar neurons with swelling formation for eventual somatic translocation, indicated with red asterisk and brackets. Through systematic microscopical examination and analysis of these series, we conclude that the AVP-ir neurons could take several routes to migrate from the SON and PVN, some of them are moving dorsomedially towards *zona incerta* (ZI), but most of them are streaming dorsolaterally towards EP, posterodorsal part of the medial amygdala (MeApd), and nucleus basalis of Meynert (B) (Fig. 1, D, green arrows).

Sagittal series of AVP immunostaining revealed massive ascending axonal routes toward the brain main conducting systems, *i.e.* fornix and stria medullaris (SF1 to Fig. 1). Numerous cell chains are discovered from this view (SF. 1 A, B. C, E. F, red arrows). AVP-ir cells with tangentially migrating morphology (*vide supra*) were found in adjacent regions to SON and PVN. Populational continuity between the SON^AVP^ and MeApd^AVP^ can be appreciated through serial section viewing (Fig. 2, from A through F).

**Fig. 2.**
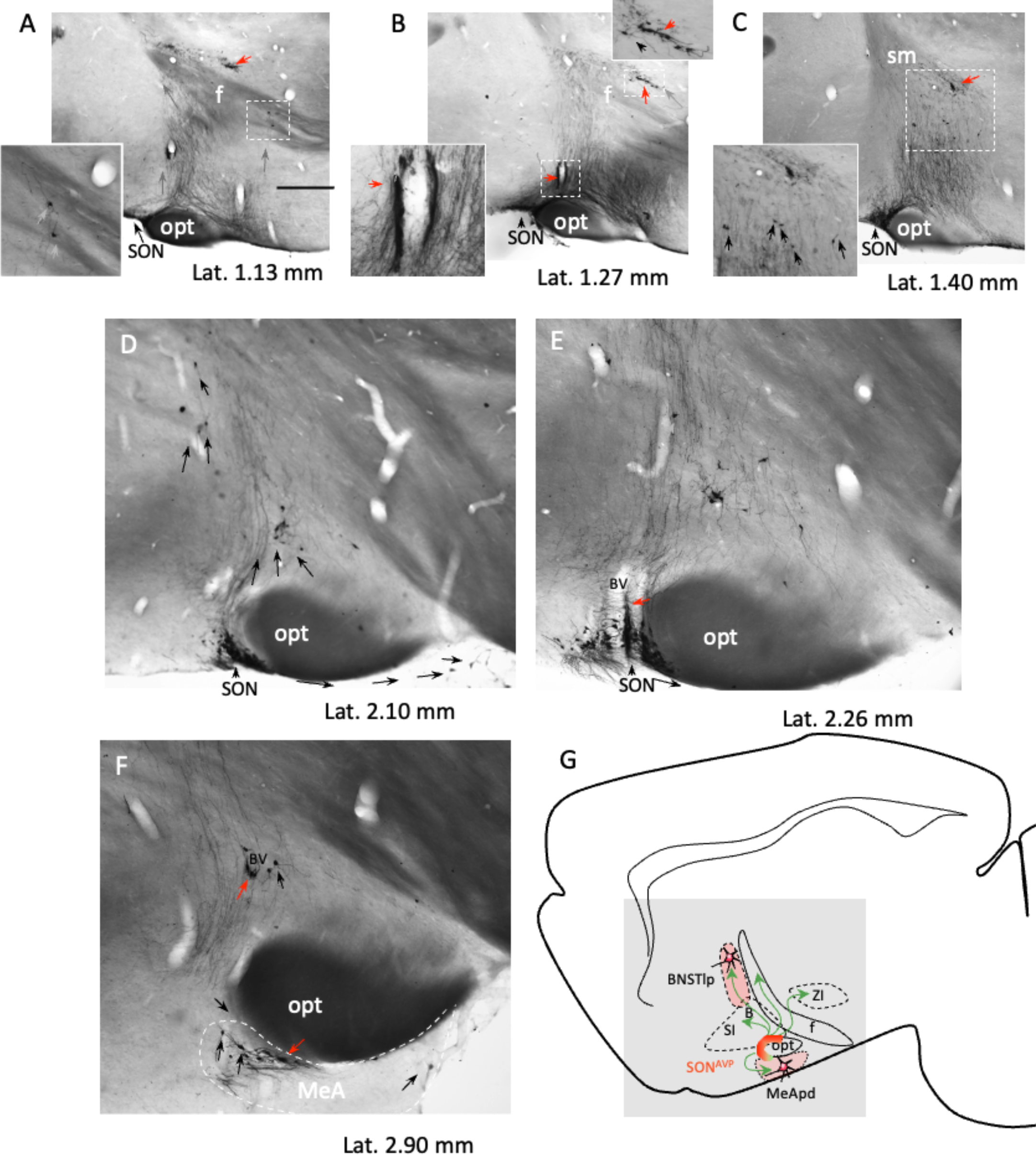
Network of AVP-ir cell chains and axon scaffolds in the adult rat hypothalamus (2/5) emerged from supraoptic nucleus (SON) - sagittal view through serial sections from an adult female Wistar rat of 4 month. A-F: AVP immunostaining at representative medio-lateral levels. Ascending axonal stream can be clearly identified. AVP-ir cell chains are indicated by red arrows. Dispersed neurons with tangential migration morphology, climbing in the AVP-ir axonal scaffold are indicated by black arrows. D, E and F show the continuity and lateral migration path of AVP-ir cells from SON to the medial amygdala alongside the optical tract (opt). G. charting symbolizing the migration routes (green arrows) visualized from sagittal series analysis. Samples were osmicated for electron microscopy study, reason that the myelinated fibers are dark. Abbreviations, B: nucleus basalis of Meynert, BSTlp: bed nucleus of stria terminalis, lateral posterior, f: fimbria, MeApd: medial amygdala posterodorsal, SI sustantia innominata, ZI: zona incerta.

Horizontal sections series of AVP-ir allowed us to appreciate that many AVP-ir neurons disperse from the SON, alongside the optic tract (opt), forming cell chains (Fig. 3, A, red arrows, note that the majority of the cell chains within the optic tract are associated with a blood vessel within the chain, indicated by red arrows). Other AVP-ir cells dispersed by tangential migration following AVP-ir axonal scaffolds (Fig. 3, A1 and A2, white arrows). Moreover, it was surprising to observe that there are at least two populations of AVP neurons withing the region of hypothalamic bed nucleus of stria terminalis (BNST-Hy). One population located in the rostral part, with small somata and weak AVP immunostaining (Fig. 3, B1 and B2, indicated by black arrowheads) and another population is posterior and at the border of the stria medullaris (sm), and the fornix (f). These latter ones are strongly immunoreactive to AVP (Fig. 3, B1 and B2, white arrows). Cell chain alongside the sm were also observed (Fig. 3, C inset, red arrows). The migratory connection between the SON and the BNST-Hy, concluded from serial section analysis, is synthesized in the diagram C of the Fig. 3. Vasopressin neuronal population’s distribution and function in the BNST-Hy has been described with ample variations among reports. For example, vasopressin in the BNST has been linked to both anxiogenic (anxiety-promoting) and anxiolytic (anxiety-reducing) effects, depending on the circumstances (Wigger, Sanchez et al. 2004). A previous study of our group reported that one subpopulation of vasopressin neurons in the BNST-Hy co-expressed VGAT, a GABAergic neuronal marker, however, a substantial portion of the cells there did not express this marker which suggest the existence of distinct subpopulations among vasopressin expressing neurons within the BNST-Hy (Zhang, Hernandez et al. 2020).

**Fig. 3.**
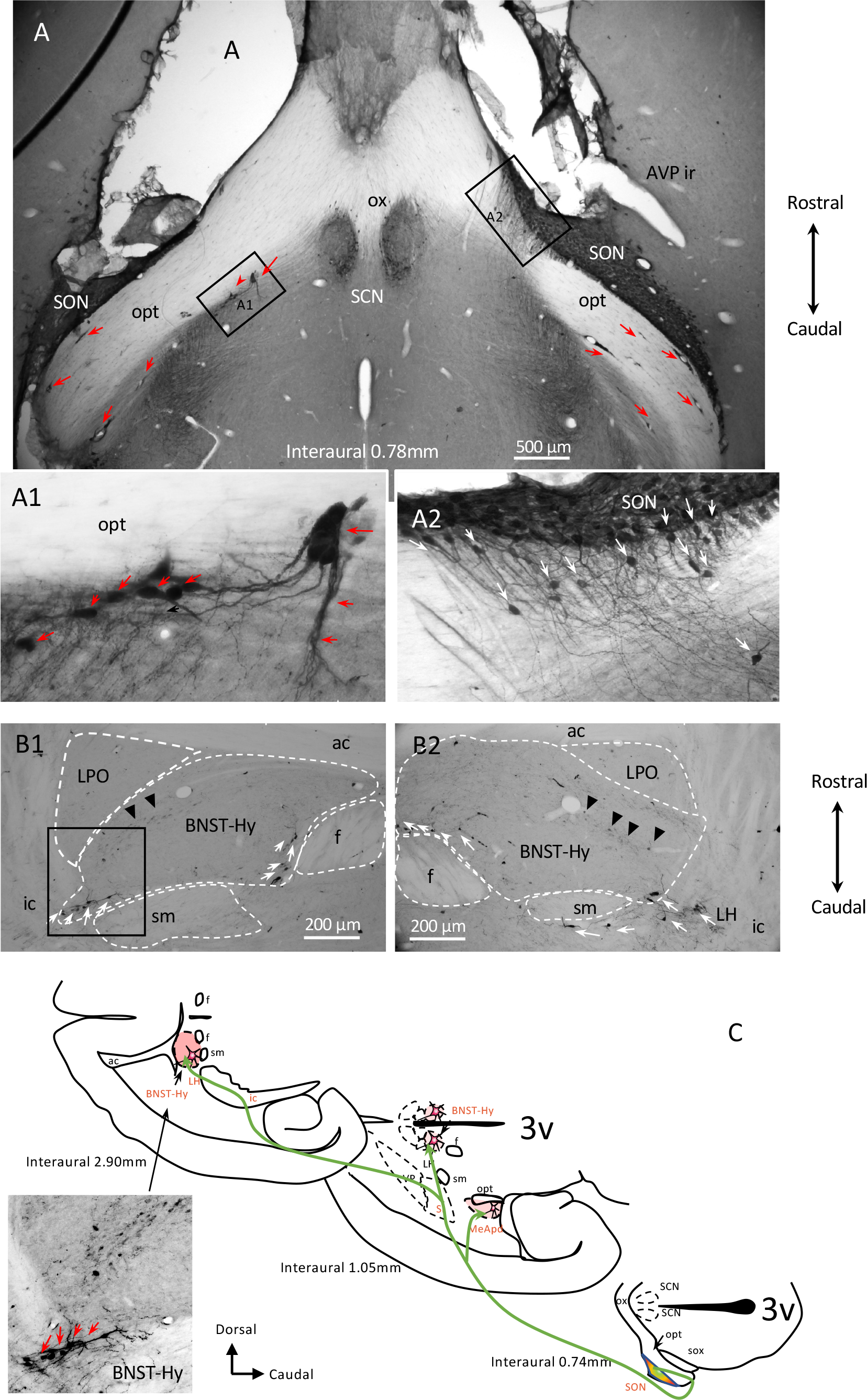
Network of AVP-ir cell chains and axon scaffolds in the adult rat hypothalamus (3/5) emerged from supraoptic nucleus (SON) - horizontal view through serial sections from an adult male Wistar rat of 4 months. As: immunostaining at the interaural level 0.78mm, where the SON and suprachiasmatic nucleus (SCN) vasopressinergic cells can be clear seen, however, with different features. While the SCN^AVP^ are well delineated, some neurons of SON^AVP^ dispersed tangentially following the axonal scaffolds (A1 and A2), mainly alongside the optic tract (opt). Numerous chains were indicated with red arrows. B1 and B2 show the tangentially migrated AVP-ir cells arriving the lateral portion of the bed nucleus of stria terminalis. One population of AVP-ir cells is located in the rostral part, with small somata and weak AVP immunostaining (indicated by black arrowheads) which can be clearly distinguished from the population migrated along the stria medullaris, and the fimbria. The latter ones are strongly immunoreactive to AVP (white arrows). C. charting symbolizing the migration routes (green arrows) from SON to medial amygdala posterodorsal division (MeApd) and bed nucleus of stria terminalis hypothalamic (BNST-Hy), passing through the sustantia innominata (SI), ventral pallidum (VP), lateral hypothalamus (LH) in 3 horizontal levels visualized by horizontal section analysis.

When the brains were sliced with a tilted angle of -30 ^o^ degree with reference to septo-temporal axis (Fig. 4, A, top right inset, red line), the immunostaining in serial sections provided a fresh view about the anatomical organization of the PVN^AVP^ in adult rat. This figure shows examples from a 12-month-old male rat. An impressive AVP-ir cell chain, alongside a main blood vessel was observed extending postero-laterally to zona incerta (ZI) (Fig. 4, red arrows). AVP-ir cells apparently dispersed from this chain, moved tangentially reaching nucleus basalis of Meynert (B) and the magnocellular nucleus of lateral hypothalamus (MCLH) (Fig. 4, A, black and white arrows). The MCLH is a group of large cells located at the lateral border of the lateral hypothalamic area at the medial border of the internal capsule of the rat (Swanson 2004). It is one of twenty six regions, zones, and nuclei constituting the motor lateral hypothalamus. Functionally it belongs to the motor lateral hypothalamus of the subcortical motor system (Swanson 2004). A bipolar AVP cell alongside a blood vessel, with axonal swelling and apparent somal translocation was indicated with a black arrow in the panel (E) of Fig. 4.

**Fig. 4.**
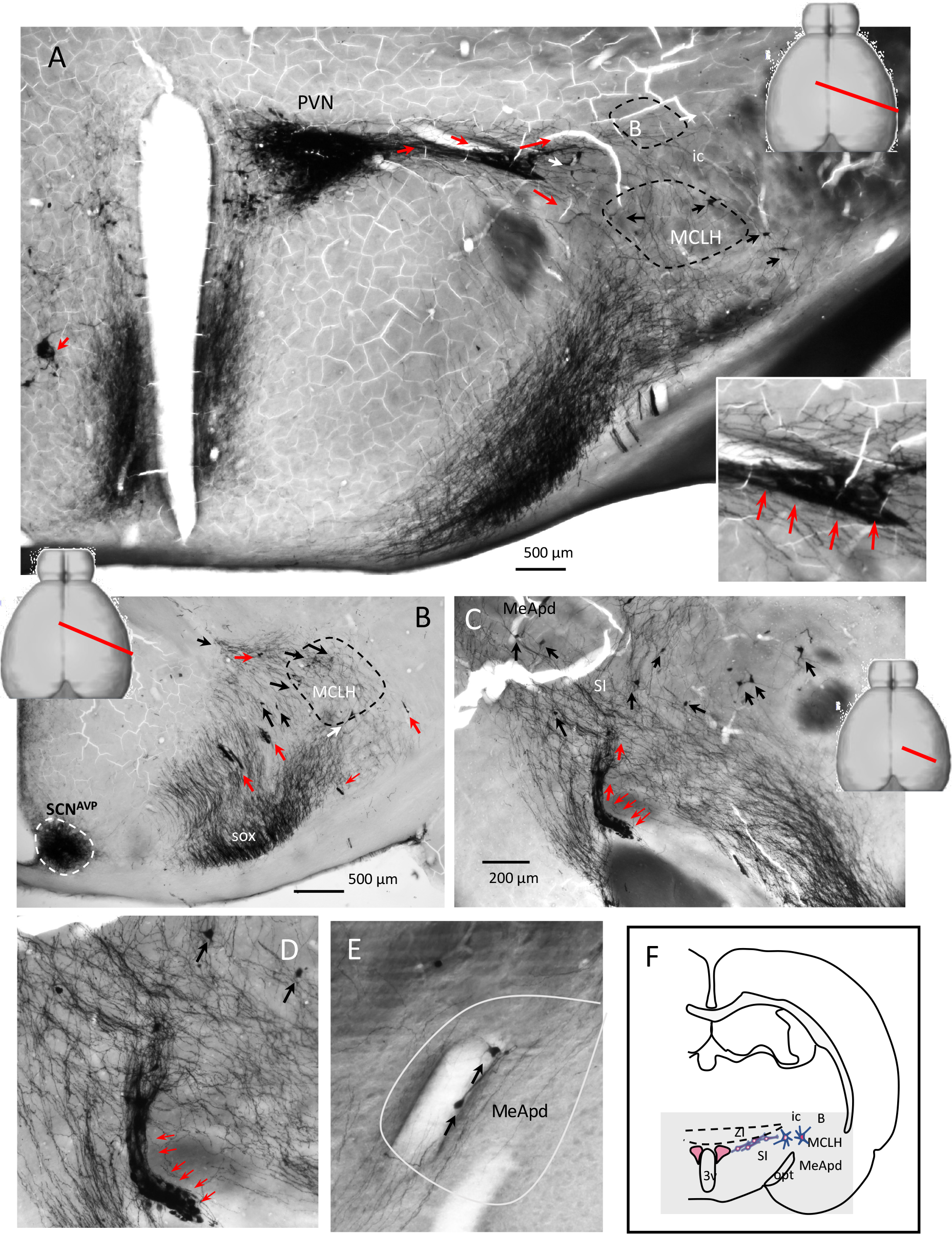
Network of AVP-ir cell chains and axon scaffolds in the adult rat hypothalamus (4/5) emerged from paraventricular (PVN) and supraoptic nucleus (SON) - septo-temporal oblique view (-30 ^o^ degree) through serial sections from a 12-month-old adult male Wistar rat. (A): AVP-ir of a septo-temporal oblique cutting plane passing through the center of PVN with -30 degrees of inclination toward temporo-caudal direction revealed thick AVP-ir cell chain emerged from PVN alongside a thick blood vessel which bifurcated to latero-dorsal (B: nucleus basalis of Meynert) and latero-ventral (MCLH: magnocellular nucleus of the lateral hypothalamus, MCLH) directions. Red arrows indicated cell chains and black arrows indicate the dispersed cells. (B): more examples of dispersed cells intermingled with axonal scaffolds. Note that the suprachiasmatic nucleus (SCN^AVP^) showed distinct feature that no cell chains or dispersed neurons were observed to emerge from there (outlined with white dashed line). (C): more examples of AVP-ir cell chains and dispersed neurons, with amplified cell chain showed in (D). Panel (E) shows tangential migrating AVP-ir cells in the medial amygdala, postero- dorsal subdivision. A bipolar AVP cell alongside a blood vessel, with axonal swelling and apparent somal translocation was indicated with a block arrow. (F): charting synthesizing the observation of this figure. Samples were osmicated for electron microscopy study, reason that the myelinated fibers are dark.

This pathway is further described in the Fig. 5, with horizontal view at interaural level 2.20 mm, where AVP-ir axon scaffolds guided cell migration can be appreciated. The PVN is shown delineated by dotted lines, and neurons migrating tangentially in a caudo-lateral direction toward the lateral hypothalamus (LH) and zona incerta (black arrows in SF. 4 to Fig. 1A). Strongly labelled AVP-ir somata were observed in the LH and ZI regions, some neurons (SF. 4 to Fig. 1 B and C, red arrows) were clustered around blood vessels (BV).

We recognize that the data presented thus far primarily *imply* postnatal migration of AVP-ir neurons. The timing of this phenomenon, whether it takes place during early stages of life or persists into adulthood, remains unanswered based on the aforementioned data.

The results mentioned above were all obtained from immunohistochemistry (IHC) mainly against vasopressin (except Fig. 1 panel A2, with IHC against neurophysin II and doublecortin) in wild-type Wistar rat brain tissue. To further address this inquiry, we aim to supplement this report with evidence obtained from a parallel study involving CRISPR/Cas9- mediated targeted insertion of the IRES-Cre transgene into the 3’ untranslated region of the AVP gene in Sprague-Dawley rats. Two adult male AVP-IRES-Cre rats were injected in the PVN with recombinant adeno-associated virus equipped by EF1a-DIO-mCherry. One month after viral vector injections, the rats were perfused and their brains were analyzed as described below.

In SF. 1 we provide examples pertain to the PVN viral vector injection in this transgenic AVP-IRES-Cre rat. In this instance, the lateral magnocellular division of the paraventricular nucleus of the hypothalamus (PVNLM, highlighted by the pink shadow region in panel A and its inset) was targeted during viral infection of the same rat line. The rat brain was fixed *one month after the viral injection*. Panel (B) and its inset illustrate the targeted site with numerous immunohistochemically labeled cells within the PVNLM^AV^ (outlined with dashed lines). The labeling was sparse yet precise, with no positive cells detected outside of the PVNLM at this medio-lateral level. Within the PVNLM^AVP^, numerous neurons exhibiting tangential migration morphology (bipolar) were observed, as indicated by red arrows. Panel (C) depicts numerous immunohistochemically labeled cells forming cellular chains within the brain’s two main conducting systems, the *fimbria* (f) and the *stria medullaris* (sm). The squared region was further magnified in (D), revealing labeled cells with tangential migrating morphology (red arrows). In panel E, migrating cells are also present in another major conducting system, the *stria terminalis* (st), with migration morphology indicated by red arrows. It is worth comparing these mCherry IHC images in transgenic rat line with SF. 1 to Fig. 1, panels A, B. and C, which are IHC against AVP in wild type rats that the very same morphological phenomena can be appreciated.

The presence of numerous neurons exhibiting tangential migration morphology within the PVNLM^AVP^ suggests active migration processes occurring in this nucleus of the adult rat brain following viral injection. SF. 1 panel (C-E) vividly illustrates the migration of labeled cells forming cellular chains within the *fornix* and *stria medullaris,* underscoring the ongoing neuronal migration processes during rat adulthood, emphasizing the dynamic nature of neuronal migration in adult rats subsequent to viral injection. It is worth noting that the *fornix*, *stria medullaris*, and *stria terminalis* constitute the main forebrain conducting systems with abundant axon bundles. The presence of neurons with migrating morphology within those conducting systems strongly suggest the transitory nature of the phenomenon, providing compelling evidence of neuron migration during adulthood.

**SF 1:**
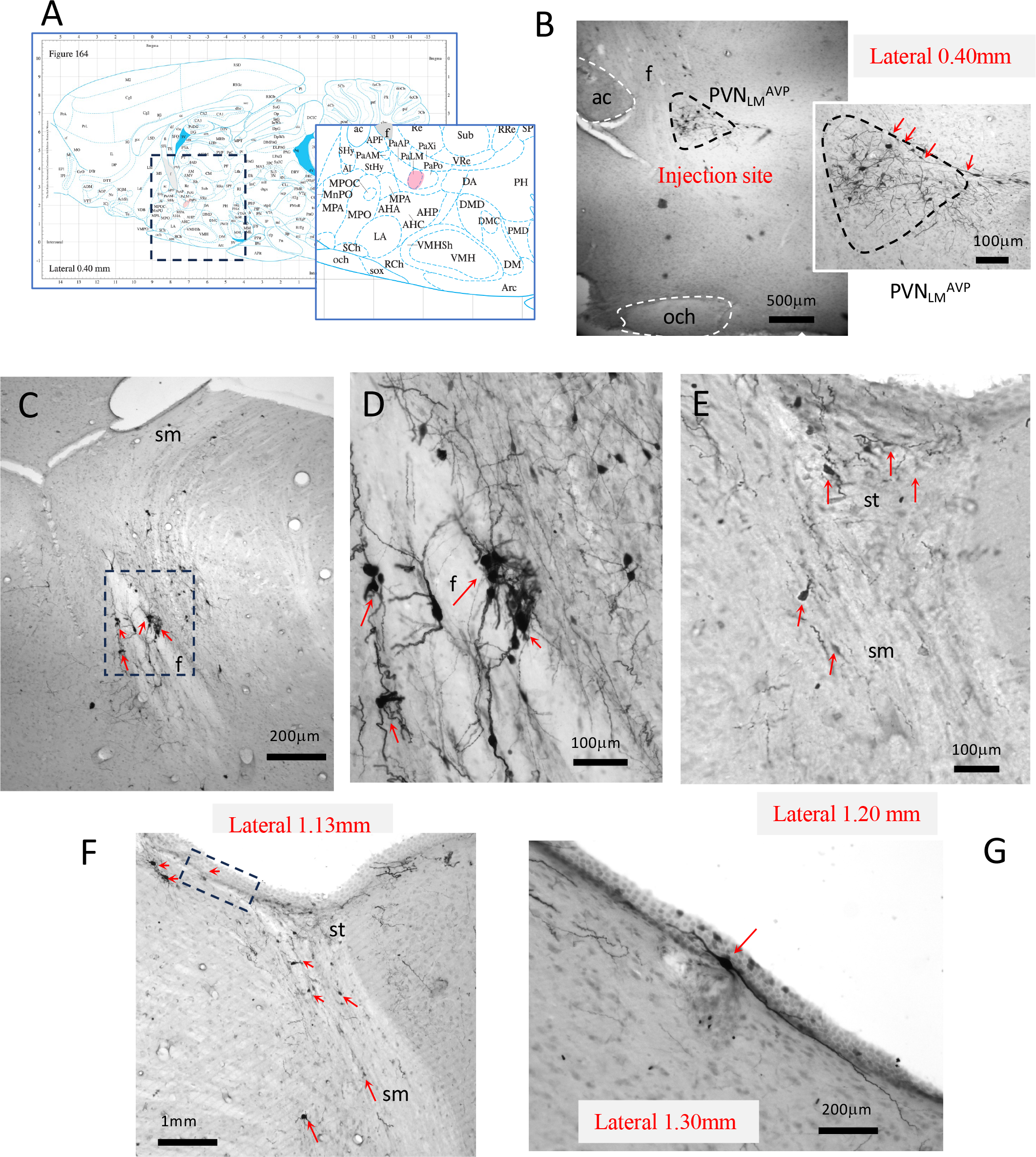
(A) and inset: Atlas sagittal plane (Paxinos and Watson, 1997) showing the location of the lateral magnocellular division of the paraventricular nucleus of the hypothalamus (PVNLM, pink shadow region), which was infected by Cre-dependent AAV equipped with mCherry in an adult male AVP-IRES-Cre knock-in rat. The rat brain was fixed after one month of the viral infection. Panel (B) and inset show the injection site with immunohistochemically (IHC) labeled cells (IHC against mCherry driven by the AVP promoter) within the PVNLM (outlined with dashed line). The labeling was sparse and precise without positive cells outside of the PVNLM. Numerous neurons with tangential migration-like morphology (bipolar) within the PVNLM were observed (indicated with red arrows). (C): photomicrograph taken from sagittal section at around lateral 1.13mm, Paxinos and Watson, 1997): numerous IHC labeled cells forming cellular chains within brain’s two main conducting systems, the fornix (f) and the stria medullaris (sm). Squared region was further enlarged in (D) showing labeled cells with tangential migrating-like morphology (red arrows). (E): photomicrograph taken from sagittal section at around lateral 1.20 mm showing putatively migrating cells present in another major conducting system, the stria terminalis (st), with migration-like morphology pointed by red arrows. (F), an additional example of the presence of the labeled cells in the sm and adjacent regions with one bipolar neuron (squared region) in close proximity to the ependyma of the third ventricle (3v) with more than 1mm of its longitudinal axis, enlarged in (G).

### Radial glial-like cells alignment reveals tangential migration paths

Glial fibrillar acidic protein (GFAP) is the principal 8-9 nm intermediate filament in mature astrocytes and radial glial cells in the central nervous system (Eng, Vanderhaeghen et al. 1971, Barry, Pakan and McDermott 2014). It is known that astrocytes, through their interaction with neuroblasts and endothelial cells, play a key role in the genesis, migration, and integration of neurons in the adult brain (Gengatharan, Bammann and Saghatelyan 2016, Bressan and Saghatelyan 2020). We used anti-GFAP IHC to explore the astrocyte distribution in the hypothalamus. Surprisingly, we found distinct patterns of long GFAP-ir processes, which seemed to form bundles representing several migration routes (Fig. 6, A, A1 and A2, indicated with white arrows). The labeling density was remarkably heterogeneous in the rat hypothalamus and the observed routes coincided with the AVP-ir cells tangential migrations routes described in Fig. 1 and its supplemental figures. Those cells appeared to resemble the tanycytes surrounding the third ventricle (3v) (Yoo and Blackshaw 2018) in the hypothalamus (Fig. 6, B), as well as the glial cells found in the adult dentate gyrus (Doetsch and Alvarez-Buylla 1996) and subgranular zone (SGZ) of the hippocampal formation (Fig. 6, C and C1). These two types of glial cells are subtypes of the radial glial cells serving for adult neuronal migration (Barry, Pakan and McDermott 2014).

**Fig. 5.**
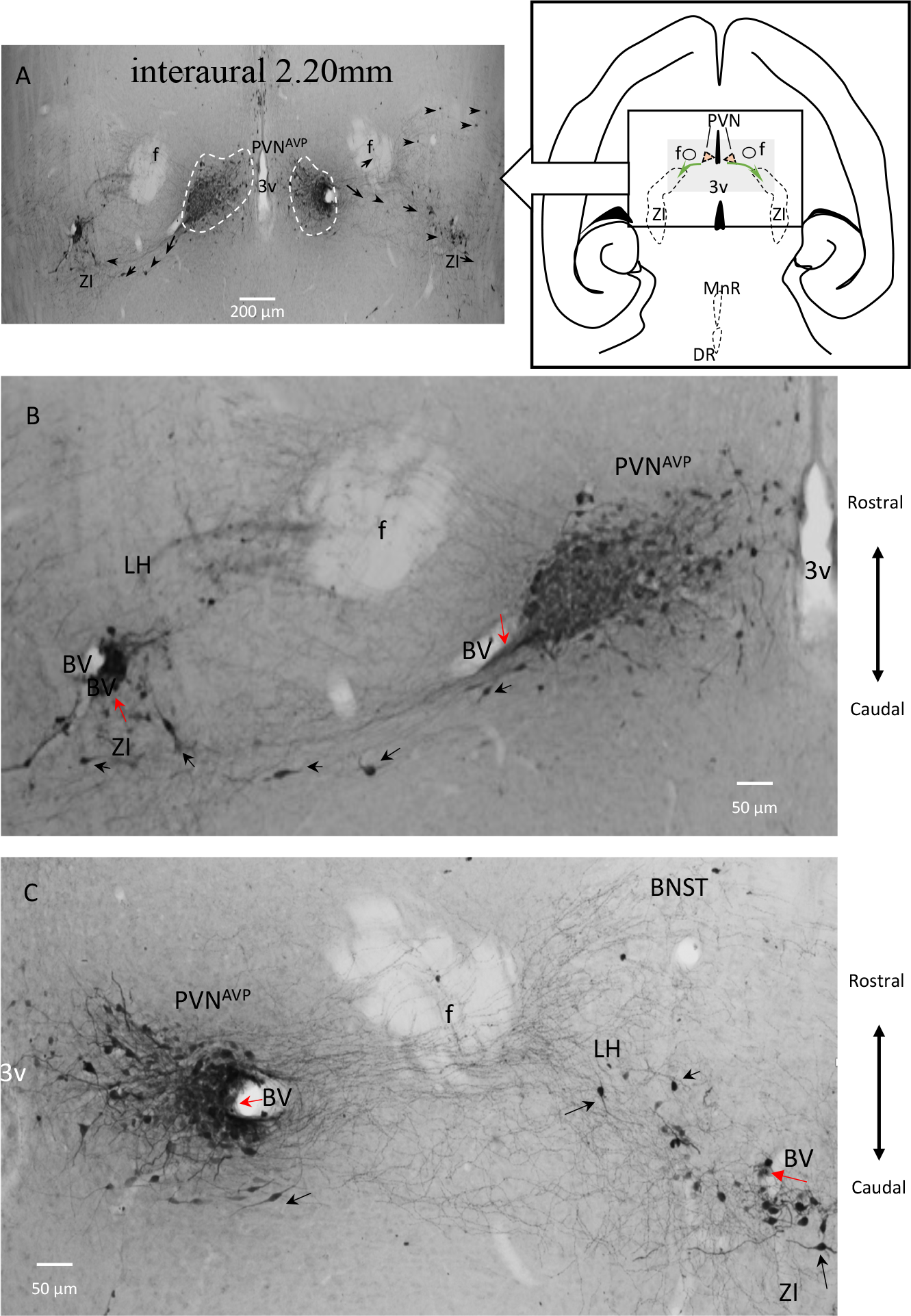
Network of AVP-ir cell chains and axon scaffolds in the adult rat hypothalamus emerged (5/5) from paraventricular nucleus (PVN) - horizontal view reveals migration route for direct connection between PVN and ZI APV-ir cells (sections from an adult male Wistar rat of 4 months). A: AVP-ir cell-chains (red arrows) and axonal scaffolds (indicate by black arrows) emerged from PVN^AVP.^ B and C are amplifications of each side of A.

**Fig. 6.**
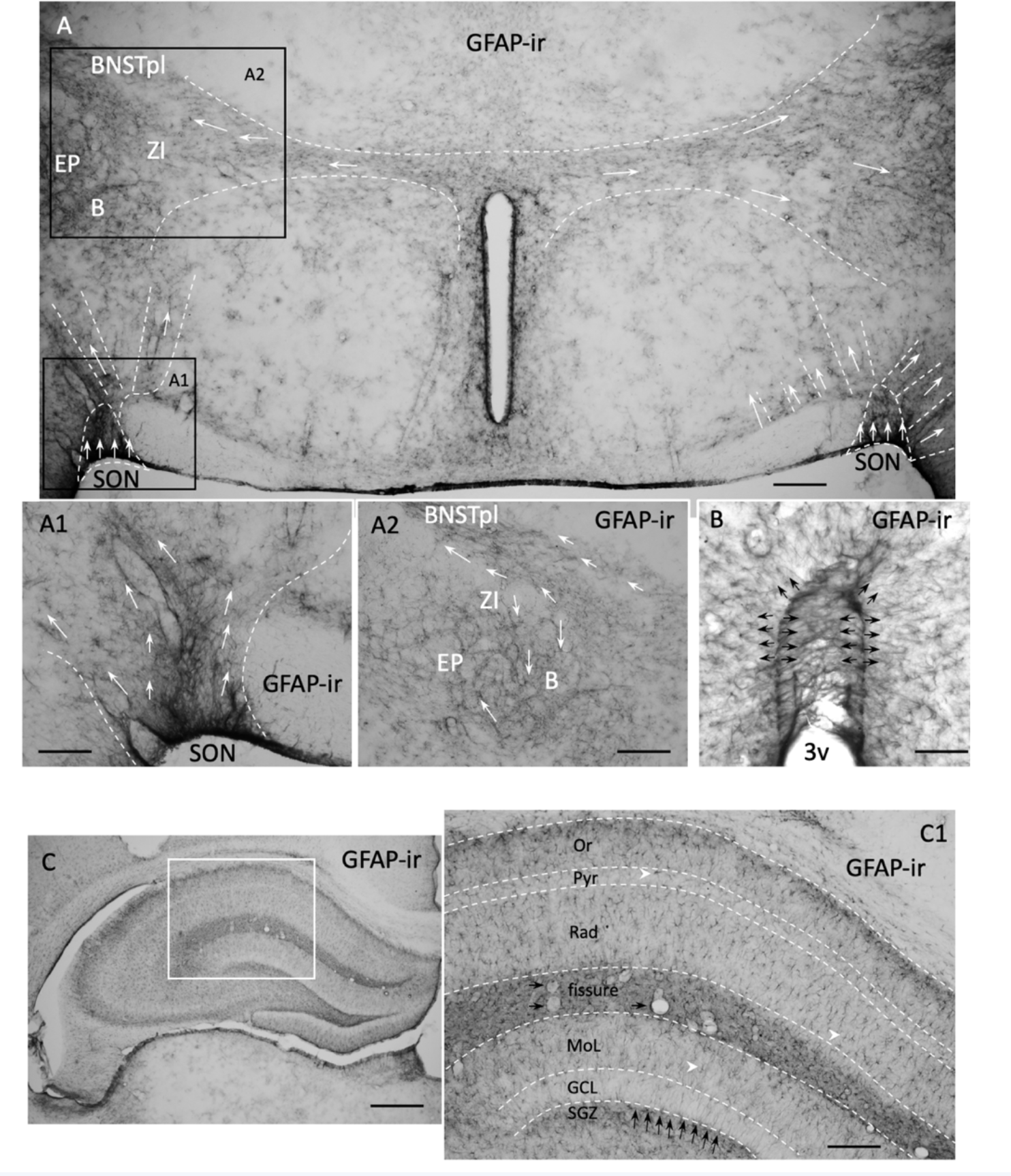
Glial fibrillar acidic protein (GFAP) immunostaining reveals a distinct feature of the GFAP- ir morphology, similar to the radial glial cell and its heterogenous distribution resembling the migration routes observed with AVP IHC in the rat hypothalamus. A, resembling patterns of tangential migration routes described above were indicated with white arrows). A1 and A2: example of distribution of the GFAP-ir cells. B. Tanycytes, a type of radial glial, in the periventricular region of the hypothalamus of the same reaction. C and inset, GFAP labeling in hippocampus. Note the similarity of the radial glial in the subgranular zone (SGZ), black arrows.

We further performed a double immunofluorescence reaction AVP/GFAP and found that all cell chains containing AVP-ir cells are closely associated with dense GFAP processes (Fig. 7 panels As, Bs and Cs). GFAP-ir long processes were seen intertwined with tangential-shaped AVP-ir cells and axons (Fig. 7, C2-C4, yellow arrows) serving as migration scaffolds for AVP- ir soma and axons (Fig.7, panels C).

**Fig. 7.**
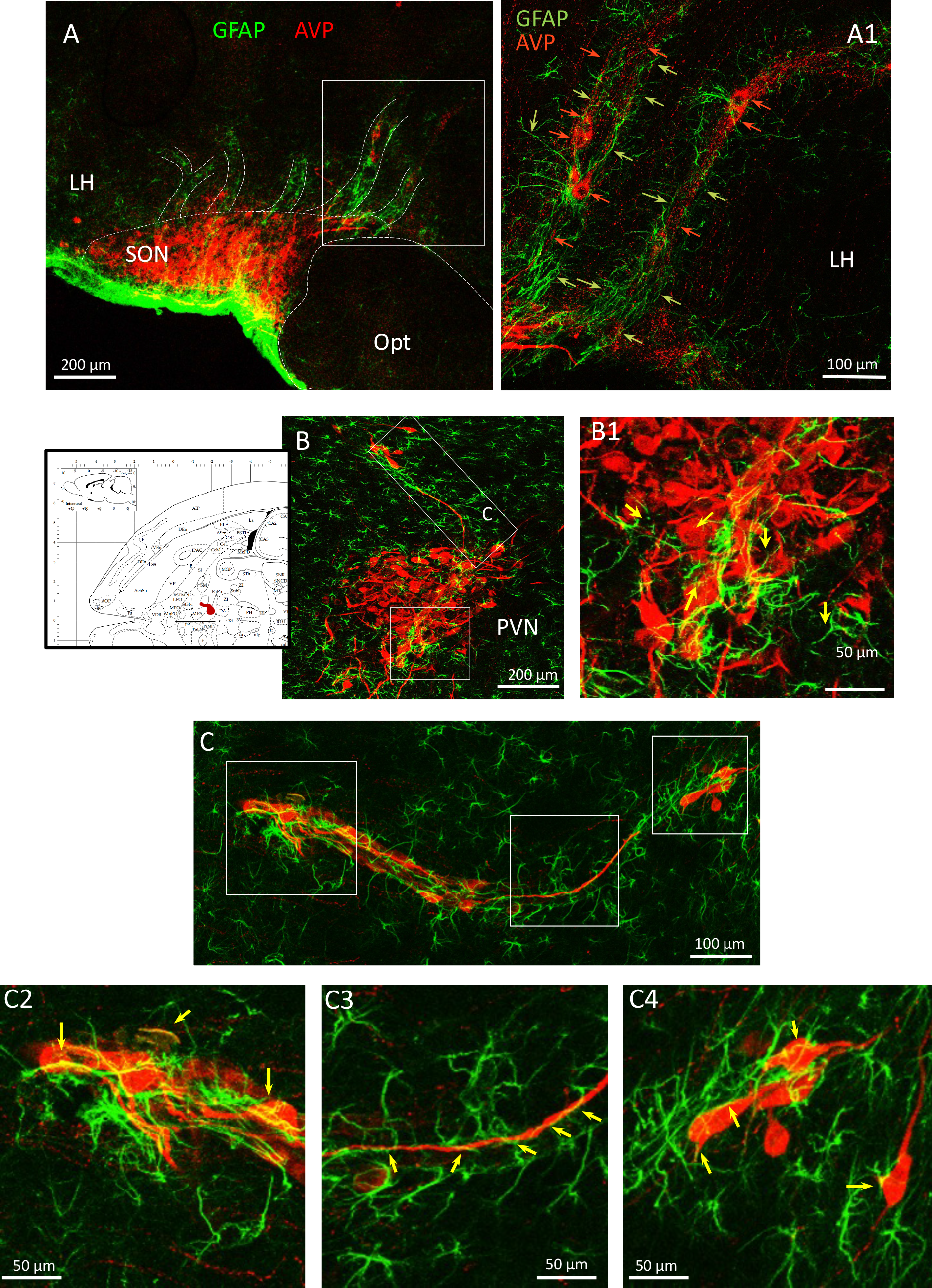
Double immunofluorescence reaction against AVP/GFAP revealed the intimate relationship between the AVP-ir dispersion, AVP neuronal chains, and the GFAP fibers. A-C, GFAP-ir tunnel-like structures containing AVP-ir cells and axons. The phenomenon of AVP-ir axons “climbing” the GFAP-ir scaffolds is indicated with yellow arrows. C2-C4 show the higher magnification of GFAP-ir long processes serving as migration scaffolds for AVP-ir soma and axons.

In order to demonstrate cell proliferation occurs within the AVP magnocelular loci, we first used immunohistochemistry against cell proliferation marker, Ki67 protein (Kee, Sivalingam et al. 2002), to test the neurogenic potential of AVP-MCN. In Fig. 8 panels A’s, two Ki67-ir nuclei adjacent to each other (A1) within a swelled weakly AVP+ cell body were shown (A and A2, dashed white line). We also used 5-bromo-2’-deoxyuridine (BrdU) injection followed by immunostaining method to evaluate cell proliferation (Gratzner 1982) in both euhydrated and chronic water-deprived (every other day, during 15 days). Panels Bs served as positive control of the procedure that show the BrdU-ir neurons within the canonical adult neurogenesis regions, B1, subventricular zone (SVZ) and B2, subgranular cell zone (SGZ) of the dentate gyrus. Figure 8 panel Cs and Ds show examples of BrdU co-labeling with neurophysin II, the carrier protein for AVP in the paraventricular (PVN) and supraoptic (SON) nuclei in both sexes, in euhydrated and water deprivation challenged rats, respectively. BrdU-ir expression was also co-assessed with microglial marker Iba1. In panels E, we show two BrdU labeled nuclei (indicated with red arrow and white arrow inside two boxes). The nucleus with indicated with white arrow has a smaller diameter and overlapped with Iba1 expression (E1 and E2), while the nucleus indicated with the red arrow has a larger diameter (7-8 μm). Panel E3 is an optical slice of 0.9 μm thickness at a difference Z level of the same cell where NPII and a branch of Iba1 fiber can be clear seen. E4 shows the relationship of IBa1 fiber with the BrdU-ir nucleus while NPII soma signal was deactivated. It seems the microglial plays a role for the newly born AVP MCN. The panel F show several complete microglial cells with Iba1-ir that its nuclear size (white arrows) is of smaller diameter (4μm) comparing with most of the BrdU labeled nuclei (7-8 μm, green arrows).

**Fig. 8.**
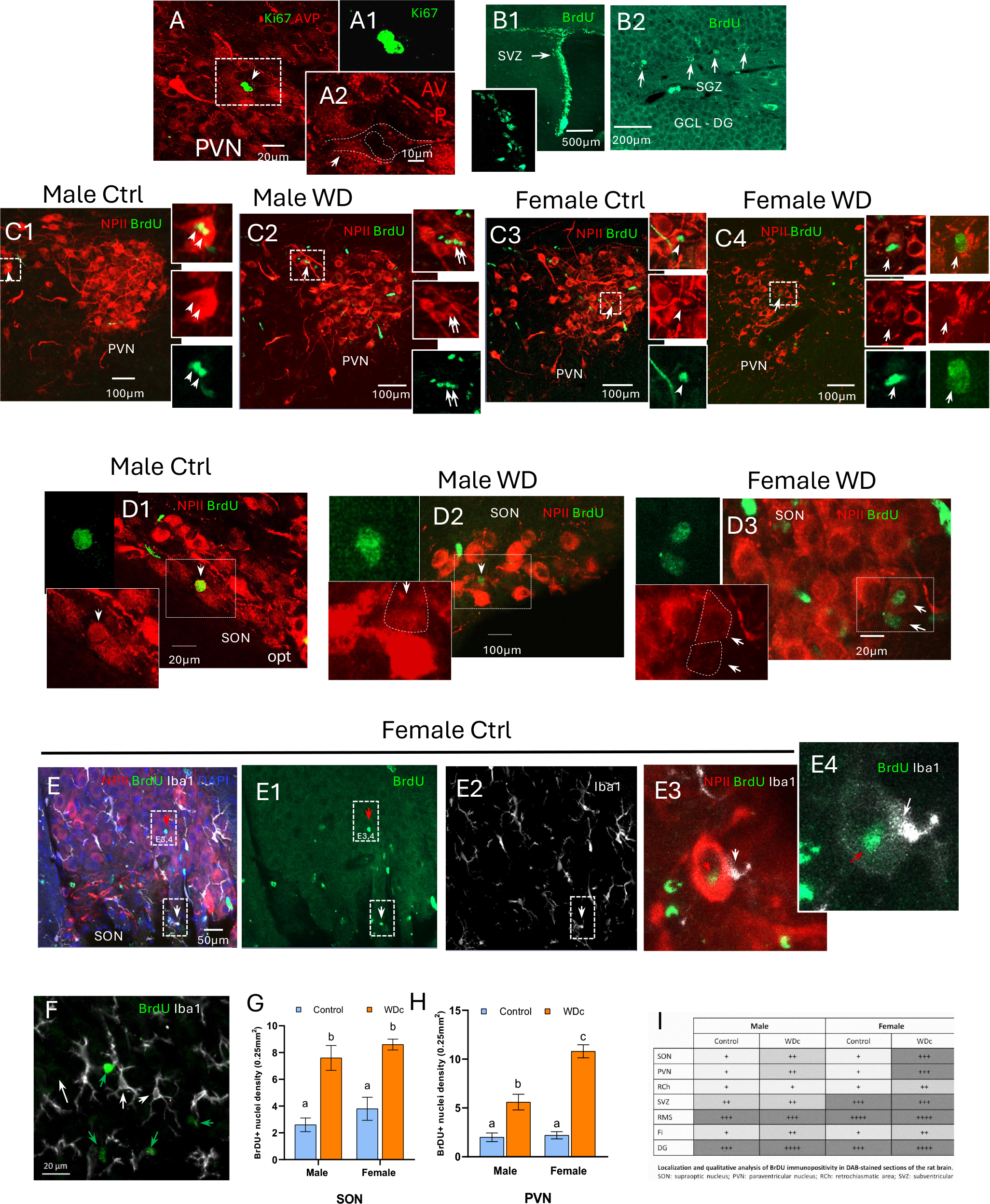
Neurogenesis occurs within the adult SON and PVN, and osmotic challenge significantly increases its rate. panels As, two Ki67-ir nuclei are adjacent to each other (A1) within a swelled weakly AVP+ cell body were shown (A and A2, dashed white line). We also used 5- bromo-2’-deoxyuridine (BrdU) injection followed by immunostaining method to evaluate cell proliferation (Gratzner 1982) in both euhydrated and chronic water-deprived (every other day, during 15 days). Panels Bs served as positive control of the procedure that show the BrdU-ir neurons within the canonical adult neurogenesis regions, B1, subventricular zone (SVZ) and B2, subgranular cell zone (SGZ) of the dentate gyrus. Figure 8 panel Cs and Ds show examples of BrdU co-labeling with neurophysin II, the carrier protein for AVP in the paraventricular (PVN) and supraoptic (SON) nuclei in both sexes, in euhydrated and water deprivation challenged rats, respectively. BrdU-ir expression was also co-assessed with microglial marker Iba1. In panels E, we show two BrdU labeled nuclei (indicated with red arrow and white arrow inside two boxes). The nucleus indicated with white arrow has a smaller diameter and overlapped with Iba1 expression (E1 and E2), while the nucleus indicated with the red arrow has a larger diameter (7-8 μm). Panel E3 is an optical slice of 0.9 μm thickness at a difference Z level of the same cell where NPII and a branch of Iba1 fiber can be clear seen. E4 shows the relationship of a microglial process making close contact to the BrdU labeled cell. For clarity, the NPII signal was deactivated in this panel. Note that it seems the microglial plays a role for the newly born AVP MCN. The panel F shows nuclear size that most of BrdU labeled nuclei are larger (7-8 μm, green arrows) than IBa1 labeled somata (white arrows) that the shade correspond to microglial nucleus is about 4 μm (NPII signal was deactivated in this panel). Panels G and H: statistical comparison of BrdU-ir nuclei in SON (G) and PVN (H), control vs chronically water-deprived subjects. Bars with different lettering indicate significant differences at p<0.05. Panel I: whole brain semi-quantitative assessment of BrdU-ir, control vs WD. Serial sagittal brain sections were analyzed and an index of positive nuclei (BrdU) in each 40x (0.45 mm) field was assessed as follows: 1-4 nuclei (+); 5-9 nuclei (++); 10-15 nuclei (+++); more than 15 nuclei (++++). SON: hypothalamic supraoptic nucleus; PVN: hypothalamic paraventricular nucleus; RCh: retro-chiasmatic area; SVZ: subventricular cell zone; RMS: rostral migratory stream; Fi: fimbria; DG: dentate gyrus.

With the observations mentioned above, we then asked if neurogenesis could be modified as an allostatic mechanism for stress adaptation. This hypothesis was assessed by exposing adult rats to intermittent water deprivation (see materials and methods for detail) and injected daily three doses of the thymidine analogue BrdU for 15 days. The same procedure of dehydration without BrdU injection was performed in order to discriminate the possibility of BrdU interference with cell proliferation. Figure 8, panels G and H: show statistical comparisons of BrdU-ir nuclei in SON (G) and PVN (H), in control vs chronically water-deprived subjects. Two-way ANOVA of SON revealed an effect of dehydration (F(1,16)=47.54, p<0.001= as the source of variation. No effect of sex (p>0.1) nor interaction between factors (p>0.5). In SON multiple comparisons of groups revealed significant differences between male controls vs male WD animals (p>0.0001), between male control vs female WD (p<0.0001) and between female control vs female WD (p<0.01). In case of PVN, differences were noted between male control vs male WD (p<0.01), between male control vs female WD (p<0.001); between male WD and female control (p<0.001). In this case, an effect of sex as a difference between male WD vs female WD (p<0.001) was detected by statistical analysis. Female control vs female WD also had differences (p<0.0001) in the number of BrdU positive nuclei. Different letters indicate differences between experimental groups. In congruence with this sexually dimorphic effect, it has been reported that estradiol can induce increased cell proliferation in arcuate and dorsomedial hypothalamus (Bless, Reddy et al. 2014). Figure 8, panel I depicts a whole brain semi-qualitative assessment of BrdU-ir, control vs WD in different brain areas where neurogenesis has been reported and the effect of WD in the BrdU-ir neuron density. As observed in the table, WD increased the number of BrdU + nuclei in SON and PVN.

### Absence of NeuN, a mature neuron marker, from SON and PVN in adult rat

NeuN is a commonly used marker for postmitotic neurons. We performed immunostaining with this marker aiming to assess the neuronal density of hypothalamus. Surprisingly, in the magnocellular PVN and SON nuclei, there was a general lack of NeuN expression (Fig. 9). In the suprachiasmatic nucleus, the ventral subdivision, which is mainly occupied by vasoactive intestinal polypeptide (VIP) - expressing neurons (Kawamoto, Nagano et al. 2003), showed a high expression of NeuN, while the dorsal part, mainly occupied by AVP-expressing neurons (Zhang, Hernandez et al. 2020), lacked NeuN expression.

**Fig. 9.**
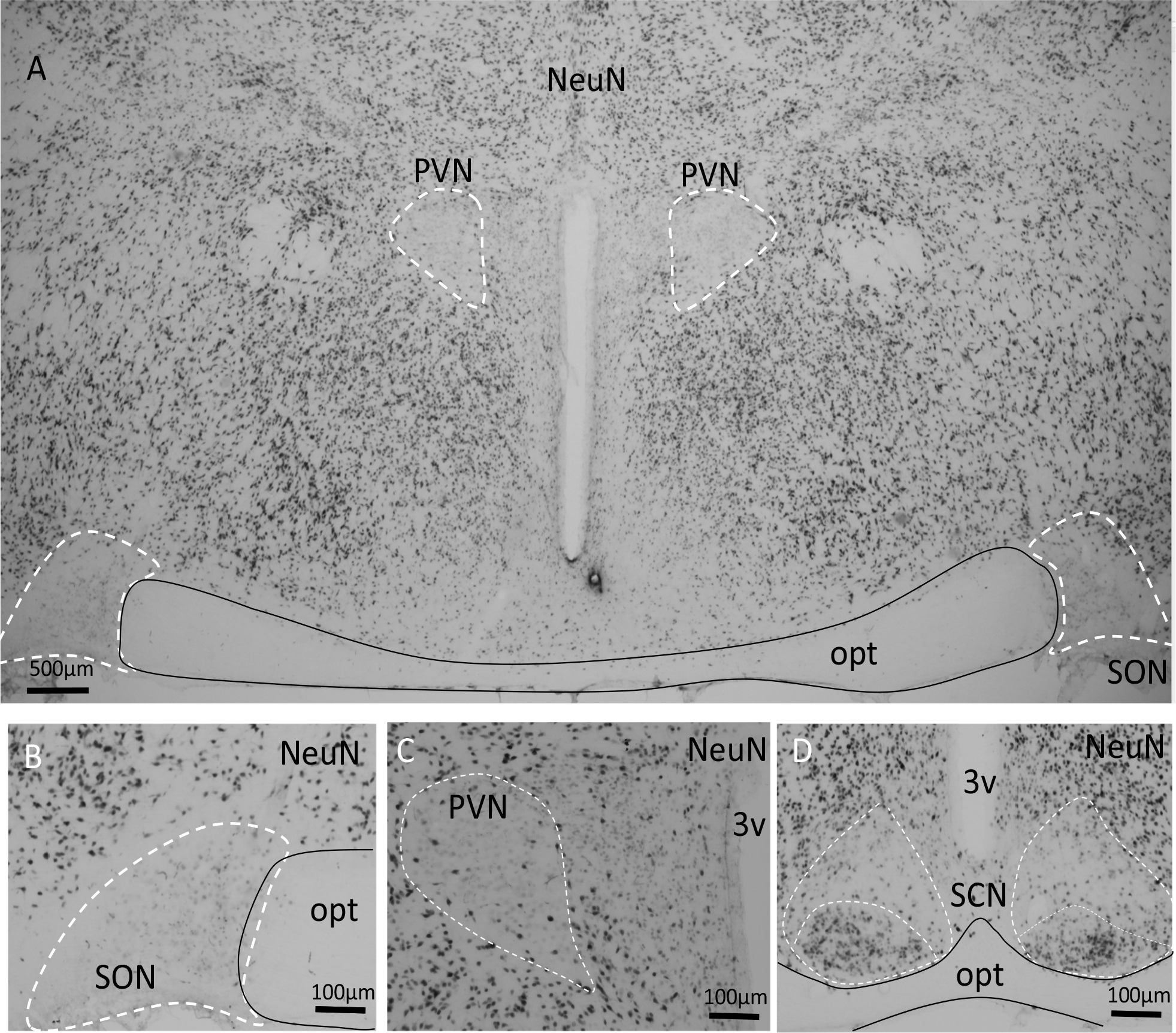
Absence of NeuN, a mature neuron marker, in the SON^AVP^ and PVN^AVP^. NeuN, a mature neuron marker, is widely expressed in the rat hypothalamus (A). However, both supraoptic (SON, A and B) and paraventricular (PVN, A and C) lack the NeuN expressing nuclei. In suprachiasmatic nucleus (D), the ventral subdivision, which is mainly occupied by vasoactive intestinal polypeptide VIP-expressing neurons, shows high expression of NeuN, and the dorsal part, which is mainly occupied by AVP-expressing neurons, lacks NeuN expression. opt: optical tract; 3v: third ventricle.

## General discussion and conclusion

During the past fifteen years, we carried out a series of systems studies on the AVP-MCN ascending projection system in adult and old rats (12-18 months), summarized in (Zhang, Hernández et al. 2021). Connections between hypothalamic AVP-MCNs and other brain areas involved in stress coping and emotional/motivational behaviors have been demonstrated, through the employment of techniques such as juxtacellular labeling and tract-tracing methods combined with brain serial sectioning in cutting planes at coronal, sagittal, septo-temporal oblique and horizontal orientations. Histochemical reaction against neurobiotin, immunohistochemistry (IHC) against antigens of interest and systematic neuroanatomical analyses were performed (Zhang and Hernandez 2013, Hernandez, Vazquez-Juarez et al. 2015, Hernandez, Hernandez et al. 2016, Zhang, Hernandez et al. 2016, Campos-Lira, Kelly et al. 2018, Hernandez-Perez, Hernandez et al. 2019, Hernandez-Perez, Hernandez et al. 2022). During these studies, a collateral observation was repeatedly made: AVP immunoreactive (AVP-ir) magnocellular neurons seemed to be dispersed from the above-mentioned loci to adjacent subcortical regions. Lateral hypothalamus (LH), zona incerta (ZI) and latero-posterior division of the bed nucleus of stria terminalis (BNSTlp) were observed as the main regions hosting these dispersed large AVP-ir neurons (DNs), but they were also seen in more distant subcortical regions, including entopeduncular nucleus (EP), nucleus basalis of Meynert (B), and postero-dorsal division of medial amygdala (MEApd). The dispersed neurons were observed to be adjacent to blood vessels and AVP-ir fibers in scaffolds, some forming cords of AVP-ir cells. These chains were visualized first with AVP IHC, with AVP-ir cell bodies within it. The phenomenon was also observed in healthy old rats (18 months). These observations collectively suggested the occurrence of adult neuronal migration, raising the questions of the mechanism of their replenishment and potential generation via adult neurogenesis within the major AVP-MCN loci (Zhou, Su et al. 2022).

In the present study, we provide evidence on the neurogenesis and migration potential of AVP-MCNs in adult rats. Through systematic neuroanatomical analysis, we conclude that an important population of AVP expressing neurons from the hypothalamic supraoptic (SON) and paraventricular (PVN) nuclei may continuously migrate to adjacent subcortical regions during adult life along streams of radial glia-like cells and processes and axon scaffolds. The recipient areas appear to include lateral hypothalamus, zona incerta, bed nucleus of stria terminalis hypothalamic lateral posterior division, nucleus basalis of Meynert, magnocellular nucleus of the lateral hypothalamus (MCLH), and entopeduncular nucleus. In adult rat brain tissue, we could frequently observe such putative tangentially migrated AVP+ cells, which appear to be following AVP+ longitudinal tunnel-like bundles and matrices, that were outlined by GFAP+ radial glial-like fibers.

The PVN and SON nuclei of the hypothalamus are known for their role in the integration of hormonal, neuroendocrine and behavioral responses as well as to be central hubs for adaptation to stress (de Wied, Diamant and Fodor 1993, Zhang, Hernandez et al. 2016, Hernandez-Perez, Hernandez et al. 2019, Hernandez-Perez, Hernandez et al. 2022). Animal studies have shown that exposure to stressors during the perinatal period leads to the increased metabolic activity of arginine-vasopressin magnocellular neurons leading to increased volume and to a robustness in their dendritic branching (Zhang, Hernandez et al. 2012).

The process that blood vessels guide the neuronal migration has been reported previously to occur with adult neuroblasts as scaffolds (Bovetti, Hsieh et al. 2007).These neuronal precursors are guided by several factors originated in the vasculature that use tyrosine kinase-associated receptors to activate migration. Examples of this are the presence of vascular brain derived neurotrophic factor, BDNF which is trapped by glial following its production by vascular cells to control the stationary/migratory phase of neuroblasts along the rostral migratory stream(Grade, Weng et al. 2013) or the vascular endothelial growth factor (VEGF), that regulates glial processes to orientate and reorganize the vascular scaffolding as well as to act as a chemoattractant to neuronal progenitors (Bozoyan, Khlghatyan and Saghatelyan 2012). These observations result of interest as there are studies showing that magnocellular neurons are able express VEFG (Wang, Li et al. 2008) and that vasopressin is able to induce the production of VEGF via the stimulation of V1aR in endothelial cells (Tahara, Tsukada et al. 2011).

We are aware about the limitation of the first finding drawn based on our anatomical observation on serial sections from wild type animals and analysis. The conclusion solely based on these data shall be only intuitive. The physiological meaning of this phenomenon, *i.e.,* adult AVP expressing MCN to disperse to other subcortical is still unclear, however, it raised important questions whether hypothalamic AVP system plays role(s) for adult brain neural protection. For instance, in a study using in situ hybridization in human tissues, Swaab and colleagues reported that the amount of AVP mRNA in hypothalamic suprachiasmatic nucleus of Alzheimer’s disease (AD) patients showed 3-fold decrease vs control (Liu, Zhou et al. 2000). In another study, the authors report that patients with AD have decreased response to hyperosmolarity challenge and are prone to dehydration (Albert, Nakra et al. 1994). A recent study (Duque, Greenwood et al. 2023, Mecawi 2023) reported intriguing multi-omic data that described the spatial transcriptome, proteome, phosphoproteome, and lipidome data from the hypothalamic supraoptic nucleus. These studies indicate that there are four groups with distinct genetic features within the SON, that are all related to AVP MCNs, offering new insight on possibilities of other AVP cell types within the SON. These cell types can be considered as candidates to study the mechanisms involved in regulating those neurons under physiological and pathological contexts (Duque, Greenwood et al. 2023).

Regarding the neurogenesis in adult hypothalamus, one of the first studies that suggested that the hypothalamus could bear neurogenic potentials was published almost twenty years ago, where the authors showed the presence of cells with immunohistochemical and electrophysiological characteristics of immature neurons in tissue cultures derived from adult rat hypothalamus (Evans, Sumners et al. 2002) and a subsequent study showed that adult rat hypothalamus cultured neurogenic precursors had the capacity to generate neuropeptide-expressing neurons (Markakis, Palmer et al. 2004). Interestingly a new population of radial glia-like neural stem cell population has been recently reported in the postnatal hypothalamus, where Irx3 and Irx5 transcription factors modulate neurogenesis of postnatally generated leptin-sensing neurons (Son, Dou et al. 2021). A recent comprehensive transcriptomic study has reported change of 2247 RNA transcripts, including Ephrin receptor 6, a gene related to adult neurogenesis, in the SON of Wistar Hannover rats’ SON AVP MCNs, after 3 days of water deprivation (Pauza, Mecawi et al. 2021).

Furthermore, the presence of DCX in AVP immunopositive neurons and the absence of the mature neuron marker NeuN suggest that a significant portion of population are neuroblasts that they have the ability to migrate as DCX is a known marker of cell proliferation as well as migration (Gleeson, Lin et al. 1999, Filipovic, Santhosh Kumar et al. 2012). The lack of expression of NeuN in the magnocellular PVN and SON nuclei and segment of the suprachiasmatic nucleus where the majority of these neurons express vasopressin, suggested a transcriptomic identity that is different to the adjacent hypothalamic nuclei. It has been reported that some neuronal populations express low levels of NeuN (Kumar and Buckmaster 2007) recently identified as an epitope of Rbfox3, an alternative splicing regulator (Kim, Adelstein et al. 2009). Interestingly, many of the regions identified as not expressing NeuN are known to exhibit postnatal neurogenesis*, i.e.* the subventricular zone (Lois and Alvarez-Buylla 1994, Lois, Garcia-Verdugo and Alvarez-Buylla 1996), the hippocampal subgranular zone of dentate gyrus (Gage, Kempermann et al. 1998), the cerebellum (Ponti, Peretto and Bonfanti 2008), and the substantia nigra (Zhao, Momma et al. 2003). A number of several other studies have shown that in local pathophysiological situations (disease/lesions) there is a reduction in the expression of NeuN without implying cellular death, but loss of immunoreactivity (Duan, Zhang et al. 2016). For instance, after axotomy of facial nerve there is a reduction of NeuN immunoactivity in facial nucleus neurons (McPhail, McBride et al. 2004); after stroke, there is a reduction in neurons expressing NeuN in the penumbra and ischemic areas (Unal-Cevik, Kilinc et al. 2004), a re- localization of NeuN to the cytoplasm of HIV infected neurons has been reported (Lucas, Calvez et al. 2014). It has also been reported that a decreased expression of NeuN immunoactivity in hippocampal neurons is linked with exposure to neurotoxins (Collombet, Masqueliez et al. 2006) and a temporary loss of NeuN immunoreactivity in hippocampal neurons after radiation (Wu, Li et al. 2010). The above-mentioned studies suggest that the non- or low- NeuN expression may be intimately linked with neuronal renewal mechanisms conserved to face brain homeostatic challenges and adaptation.

## Materials and methods

### 2.1 Animals

This study was performed using thirty-four Wistar rats. All animal procedures were approved by the local bioethical committee. Rats were housed in plexiglass cages (2-3 per cage) attached to an air recycler device, with controlled humidity, temperature (22-25°C), and filtered air ventilation, under controlled illumination (12:12 h light/dark photocycles). Food and water were provided *ad libitum* unless indicated otherwise.

### 2.2 Experimental design

The study was divided in two parts. In the first part, systematic analysis and documentation of the patterns that some AVP neurons dispersed from SON and PVN were made by anatomical examination of permanently mounted *serial sections* samples with *coronal, sagittal, horizontal and septo-temporal oblique* orientations, which were accumulated from our previous studies. The ethical statements, animal experiment licenses and experimental details of those studies can be found elsewhere (Zhang and Hernandez 2013, Irles, Nava-Kopp et al. 2014, Hernandez, Vazquez-Juarez et al. 2015, Hernandez, Hernandez et al. 2016, Zhang, Hernandez et al. 2016, Campos-Lira, Kelly et al. 2018, Hernandez-Perez, Hernandez et al. 2019, Hernandez-Perez, Hernandez et al. 2022).

The second part concerned neurogenesis detection and demonstration using 5′- bromo-2′-deoxyuridine (BrdU) protocol (Zhang, Hernandez et al. 2014) and immunohistochemistry (IHC) against cell proliferation markers BrdU and Ki67, as well as migrating neuroblast marker doublecortin (DCX), and their relationship with GFAP and AVP expression. This part was performed under the license FM-CIE-079-2020.

### 2.3 Chemicals

Chemicals were obtained from Sigma–Aldrich, St. Louis, MO, USA, if not indicated otherwise. Sources of primary antibodies and their dilutions are depicted in Table 1.

### 2.4 Experimental treatment of water deprivation (WD) and BrdU injection

Twenty-four rats (250+/-20g, 12 per each sex) were used in this experiment which consisted of two protocols. Rats were divided into two groups: controls, n=6 x 2 (male and female) and water deprivation WD, n=6 x 2 (sexes). WD subjects were restricted from water intake every other 24 hrs. For protocol A, n=4, intraperitoneal injection of the thymidine analog 5- bromo-2-deoxiurydine (BrdU, Sigma Aldrich) diluted in NaCl 0.9% (10 mg/ml) was given to each experimental subject for 14 days, with daily doses of 50 mg/kg body weight fractionated in 3 injections per day. Protocol B consisted in the same control and water deprivation treatment but without BrdU injection (n=2 per group, N=8) for Ki-67, GFAP, and DCX immunolabeling vs. AVP-ir.

### 2.5 Immunohistochemistry

Rats were deeply anesthetized with an overdose of sodium pentobarbital (63mg/kg, Sedalpharma, Mexico) and perfused trans-aortically with 0.9% saline followed by cold fixative containing 4% of paraformaldehyde in 0.1 M sodium phosphate buffer (PB, pH 7.4) plus 15% v/v saturated picric acid for 15 min (referred as Somogyi fixative (Somogyi and Takagi 1982)). Brains were immediately removed, blocked, then thoroughly rinsed with PB. Immunohistochemical reactions (IHC) were performed the same day of perfusion/fixation and sectioning (N. B. this timing scheme has given us a remarkably better AVP labeling) using the immunoperoxidase (for single antigen labeling) and immunofluorescence (multi-antigen labeling) methods. The IHC procedure of previous studies can be found elsewhere (Zhang and Hernandez 2013, Irles, Nava-Kopp et al. 2014, Hernandez, Vazquez-Juarez et al. 2015, Hernandez, Hernandez et al. 2016, Zhang, Hernandez et al. 2016, Campos-Lira, Kelly et al. 2018, Hernandez-Perez, Hernandez et al. 2019, Hernandez-Perez, Hernandez et al. 2022). Briefly, brains were sectioned using a Leica VT 1000S vibratome, at 70 µm thickness, soon after perfusion/fixation (15 min) and thoroughly rinsed (until the yellow color of picric acid was cleared). The following sectioning planes were used: sagittal (n = 3), coronal (n = 3), semi-horizontal (30° to the horizontal plane, n = 2) and septo-temporal (between coronal and sagittal planes, 45° to both planes, n = 2). Freshly-cut freely-floating sections (1 in 4) from different cutting planes were blocked with 20% normal donkey serum (NDS) in Tris-buffered (0.05 M, pH 7.4) saline (0.9%) plus 0.3% of Triton X-100 (TBST) for 1 h at room temperature and incubated with the primary antibodies listed in the table 1 in TBST plus 1% NDS over two nights at 4°C with gentle shaking. For immunoperoxidase reaction, conventional procedure were followed (detailed description can be found (Zhang and Hernandez 2013, Irles, Nava-Kopp et al. 2014, Hernandez, Vazquez-Juarez et al. 2015, Hernandez, Hernandez et al. 2016, Zhang, Hernandez et al. 2016, Campos-Lira, Kelly et al. 2018, Hernandez-Perez, Hernandez et al. 2019, Hernandez-Perez, Hernandez et al. 2022). For BrdU-IHC, sections were pretreated by incubating sections in 2N HCl for 30 minutes at 37°C for DNA denaturalization. After this step, sections were rinsed twice with 0.1 M borate buffer (pH 8.5) followed by a rinse with phosphate buffer (Zhang, Guadarrama et al. 2006).

### 2.6 Transmission electron microscopy

Technical details of electron microscopy photomicrograph displayed in the inset of figure 1 were reported elsewhere(Zhang and Hernandez 2013).

## Supplemental Methods

The transgenic rat strain, AVP-IRES2-Cre knock-in rats, was generated by CRISPR/Cas9-mediated targeted insertion of the IRES2-Cre transgene in the 3’ untranslated region of the AVP gene in Sprague-Dawley rats (Krabichler, Lefevre et al. 2023). This strain allows us to specifically target distinct AVP neurons via stereotaxic injections of Cre-dependent viral vectors, to express any genes of interest. Two adult AVP-IRES2-Cre knock-in transgenic rats (one female and one male) were injected with viral vector one month before the perfusion-fixation. Fixed brain tissues were sliced in sagittal plane with a Leica vibratome (VT1000) and serial sections were processed, immunostained with a goat anti mCherry antibody (Goat tdTomato Polyclonal Antibody-AEG42798.1, MBS448092, MYBIORESOURCE, San Diego, CA). This antibody (MBS448092) is specific for tdTomato and mCherry proteins). Samples were incubated with primary antibody (1:2000) over two night, at 4°C, with gentle shacking and then incubated with secondary antibody, rabbit anti-Goat IgG Antibody (H+L), biotinylated (BA-5000-1.5), 1:500 (Vector Laboratories) over night, at 4°C (this procedure produces good ultrastructure, if examined under electron microscope). Finishing the secondary antibody incubation, samples were incubated with avidin-biotin-peroxidase complex (ABC) in immunoperoxidase techniques (Vector Laboratories). DAB (3,3’-Diaminobenzidine, 0.05% in 0.1M phosphate buffer, pH7.4) was oxidized by hydrogen peroxide in a reaction typically catalyzed by horseradish peroxidase (HRP) which was linked to the B component of the ABC complex. The oxidized DAB forms a brown precipitate, at the location of the HRP, linked to the presence of mCherry, which was linked the vasopressin promotor. Samples were then examined under light microscopy.

## Acknowledgement

The research leading to these results received funding from the National Autonomous University of Mexico (UNAM), under Grant Agreements UNAM-PAPIIT-IG200121 and from Mexican Nacional Council for Humanity, Science and Technology (CONAHCYT), under Grant Agreement CF-2023-243. The investigators of this study have been supported by the following fellowships: sabbatical fellowships from the PASPA program of the Dirección General de Personal Académico (DGAPA) of the UNAM (LZ, VSH) and CONAHCYT (LZ) for a sabbatical research year hosted by Dr. Lee E. Eiden in the Section on Molecular Neuroscience of the NIMH-IRP, NIH, USA; UNAM-DGAPA-POSDOC program of DGAPA-UNAM (MZ) and Fulbright-García Robles Fellowship (VSH). We thank Dr. Lee E. Eiden for his constant supports and for careful and critical reading and correction of an early version of this manuscript. We are grateful to Professors Peter Somogyi, Ruud Buijs, David Murphy, Dr. Sunny Jiang for in-person examination of samples and discussing the findings of this study together with us. Our thanks also go to Dr. Marina Eliava, Alan Kania, Alicia Nava-Kopp, Erika Vazquez-Juárez that some samples examined in this study were prepared with their assistance.

We wish to dedicate this original research paper to the memory of Dr. Harold (Hal) Gainer, whose passion for neuroscience and unwavering dedication to advancing the magnocellular neurosecretory research field continue to inspire us all, and his generosity in mentoring and supporting the younger generations enriched our lives.

## References

1. Albert, S. G., B. R. Nakra, G. T. Grossberg and E. R. Caminal (1994). “Drinking behavior and vasopressin responses to hyperosmolality in Alzheimer’s disease.” Int Psychogeriatr 6(1): 79–86.

2. Altman, J. (1962). “Are new neurons formed in the brains of adult mammals?” Science 135(3509): 1127–1128.

3. Altman, J. (1963). “Autoradiographic investigation of cell proliferation in the brains of rats and cats.” Anat Rec 145: 573–591.

4. Altman, J. and G. D. Das (1966). “Autoradiographic and histological studies of postnatal neurogenesis. I. A longitudinal investigation of the kinetics, migration and transformation of cells incorporating tritiated thymidine in neonate rats, with special reference to postnatal neurogenesis in some brain regions.” J Comp Neurol 126(3): 337–389.

5. Armstrong, W. (2004). Hypothalamic supraoptic and paraventricular nuclei. The Rat Nervous System,. G. Paxinos. Amsterdam, Elsevier: 369–388.

6. Bargmann, W. and E. Scharrer (1951). “The site of origin of the hormones of the posterior pituitary.” Am Sci 39(2): 255–259.

7. Barry, D. S., J. M. Pakan and K. W. McDermott (2014). “Radial glial cells: key organisers in CNS development.” Int J Biochem Cell Biol 46: 76–79.

8. Bless, E. P., T. Reddy, K. D. Acharya, B. S. Beltz and M. J. Tetel (2014). “Oestradiol and diet modulate energy homeostasis and hypothalamic neurogenesis in the adult female mouse.” J Neuroendocrinol 26(11): 805–816.

9. Bovetti, S., Y. C. Hsieh, P. Bovolin, I. Perroteau, T. Kazunori and A. C. Puche (2007). “Blood vessels form a scaffold for neuroblast migration in the adult olfactory bulb.” J Neurosci 27(22): 5976–5980.

10. Bozoyan, L., J. Khlghatyan and A. Saghatelyan (2012). “Astrocytes control the development of the migration-promoting vasculature scaffold in the postnatal brain via VEGF signaling.” J Neurosci 32(5): 1687–1704.

11. Bressan, C. and A. Saghatelyan (2020). “Intrinsic Mechanisms Regulating Neuronal Migration in the Postnatal Brain.” Front Cell Neurosci 14: 620379.

12. Buijs, R. M., C. W. Pool, J. J. Van Heerikhuize, A. A. Sluiter, P. J. Van de Sluis, M. Ramkema, T. P. Van der Woude and E. Van der Beek (1989). “Antibodies to small transmitter molecules and peptides: production and application of antibodies to dopamine, serotonin, GABA, vasopressin, vasoactive intestinal peptide, neuropeptide Y, somatostatin and substance P.” Biomedical Research 10: 213–221.

13. Campos-Lira, E., L. Kelly, M. Seifi, T. Jackson, T. Giesecke, K. Mutig, T. A. Koshimizu, V. S. Hernandez, L. Zhang and J. D. Swinny (2018). “Dynamic Modulation of Mouse Locus Coeruleus Neurons by Vasopressin 1a and 1b Receptors.” Front Neurosci 12: 919.

14. Cheng, M. F. (2013). “Hypothalamic neurogenesis in the adult brain.” Front Neuroendocrinol 34(3): 167–178.

15. Collombet, J. M., C. Masqueliez, E. Four, M. F. Burckhart, D. Bernabe, D. Baubichon and G. Lallement (2006). “Early reduction of NeuN antigenicity induced by soman poisoning in mice can be used to predict delayed neuronal degeneration in the hippocampus.” Neurosci Lett 398(3): 337–342.

16. Cunningham, E. T., Jr. and P. E. Sawchenko (1991). “Reflex control of magnocellular vasopressin and oxytocin secretion.” Trends Neurosci 14(9): 406–411.

17. Das, G. D. and J. Altman (1970). “Postnatal neurogenesis in the caudate nucleus and nucleus accumbens septi in the rat.” Brain Res 21(1): 122–127.

18. de Wied, D., M. Diamant and M. Fodor (1993). “Central nervous system effects of the neurohypophyseal hormones and related peptides.” Front Neuroendocrinol 14(4): 251–302.

19. Doetsch, F. and A. Alvarez-Buylla (1996). “Network of tangential pathways for neuronal migration in adult mammalian brain.” Proc Natl Acad Sci U S A 93(25): 14895–14900.

20. Duan, W., Y. P. Zhang, Z. Hou, C. Huang, H. Zhu, C. Q. Zhang and Q. Yin (2016). “Novel Insights into NeuN: from Neuronal Marker to Splicing Regulator.” Mol Neurobiol 53(3): 1637–1647.

21. Duque, V., M. Greenwood, M. D., . A.S., T. Camilo, A. Pauza and B. Gillard (2023). Single-nucleus RNA-Seq revealed the core genes related to the magnocellular neurons activity. Pan-American Physiological Sciences 2023, Puerto Varas, Chile.

22. Eng, L. F., J. J. Vanderhaeghen, A. Bignami and B. Gerstl (1971). “An acidic protein isolated from fibrous astrocytes.” Brain Res 28(2): 351–354.

23. Ernst, A., K. Alkass, S. Bernard, M. Salehpour, S. Perl, J. Tisdale, G. Possnert, H. Druid and J. Frisen (2014). “Neurogenesis in the striatum of the adult human brain.” Cell 156(5): 1072–1083.

24. Evans, J., C. Sumners, J. Moore, M. J. Huentelman, J. Deng, C. H. Gelband and G. Shaw (2002). “Characterization of mitotic neurons derived from adult rat hypothalamus and brain stem.” J Neurophysiol 87(2): 1076–1085.

25. Falke, N. (1991). “Modulation of oxytocin and vasopressin release at the level of the neurohypophysis.” Prog Neurobiol 36(6): 465–484.

26. Filipovic, R., S. Santhosh Kumar, C. Fiondella and J. Loturco (2012). “Increasing doublecortin expression promotes migration of human embryonic stem cell-derived neurons.” Stem Cells 30(9): 1852–1862.

27. Gage, F. H., G. Kempermann, T. D. Palmer, D. A. Peterson and J. Ray (1998). “Multipotent progenitor cells in the adult dentate gyrus.” J Neurobiol 36(2): 249–266.

28. Gengatharan, A., R. R. Bammann and A. Saghatelyan (2016). “The Role of Astrocytes in the Generation, Migration, and Integration of New Neurons in the Adult Olfactory Bulb.” Front Neurosci 10: 149.

29. Gleeson, J. G., P. T. Lin, L. A. Flanagan and C. A. Walsh (1999). “Doublecortin is a microtubule-associated protein and is expressed widely by migrating neurons.” Neuron 23(2): 257–271.

30. Grade, S., Y. C. Weng, M. Snapyan, J. Kriz, J. O. Malva and A. Saghatelyan (2013). “Brain-derived neurotrophic factor promotes vasculature-associated migration of neuronal precursors toward the ischemic striatum.” PLoS One 8(1): e55039.

31. Gratzner, H. G. (1982). “Monoclonal antibody to 5-bromo- and 5-iododeoxyuridine: A new reagent for detection of DNA replication.” Science 218(4571): 474–475.

32. Hatton, G. I. (1990). “Emerging concepts of structure-function dynamics in adult brain: the hypothalamo-neurohypophysial system.” Prog Neurobiol 34(6): 437–504.

33. Hernandez, V. S., O. R. Hernandez, M. Perez de la Mora, M. J. Gomora, K. Fuxe, L. E. Eiden and L. Zhang (2016). “Hypothalamic Vasopressinergic Projections Innervate Central Amygdala GABAergic Neurons: Implications for Anxiety and Stress Coping.” Front Neural Circuits 10: 92.

34. Hernandez, V. S., E. Vazquez-Juarez, M. M. Marquez, F. Jauregui-Huerta, R. A. Barrio and L. Zhang (2015). “Extra-neurohypophyseal axonal projections from individual vasopressin-containing magnocellular neurons in rat hypothalamus.” Front Neuroanat 9: 130.

35. Hernandez-Perez, O. R., V. S. Hernandez, A. T. Nava-Kopp, R. A. Barrio, M. Seifi, J. D. Swinny, L. E. Eiden and L. Zhang (2019). “A Synaptically Connected Hypothalamic Magnocellular Vasopressin-Locus Coeruleus Neuronal Circuit and Its Plasticity in Response to Emotional and Physiological Stress.” Front Neurosci 13: 196.

36. Hernandez-Perez, O. R., V. S. Hernandez, M. A. Zetter, L. E. Eiden and L. Zhang (2022). “Nucleus of the lateral olfactory tract: A hub linking the water homeostasis-associated supraoptic nucleus-arginine vasopressin circuit and neocortical regions to promote social behavior under osmotic challenge.” J Neuroendocrinol: e13202.

37. Irles, C., A. T. Nava-Kopp, J. Moran and L. Zhang (2014). “Neonatal maternal separation up-regulates protein signalling for cell survival in rat hypothalamus.” Stress 17(3): 275–284.

38. Jurkowski, M. P., L. Bettio, K. W. E, A. Patten, S. Y. Yau and J. Gil-Mohapel (2020). “Beyond the Hippocampus and the SVZ: Adult Neurogenesis Throughout the Brain.” Front Cell Neurosci 14: 576444.

39. Kaneko, N., M. Sawada and K. Sawamoto (2017). “Mechanisms of neuronal migration in the adult brain.” J Neurochem 141(6): 835-847.

40. Kawamoto, K., M. Nagano, F. Kanda, K. Chihara, Y. Shigeyoshi and H. Okamura (2003). “Two types of VIP neuronal components in rat suprachiasmatic nucleus.” J Neurosci Res 74(6): 852–857.

41. Kee, N., S. Sivalingam, R. Boonstra and J. M. Wojtowicz (2002). “The utility of Ki-67 and BrdU as proliferative markers of adult neurogenesis.” J Neurosci Methods 115(1): 97–105.

42. Kokoeva, M. V., H. Yin and J. S. Flier (2005). “Neurogenesis in the hypothalamus of adult mice: potential role in energy balance.” Science 310(5748): 679–683.

43. Kokoeva, M. V., H. Yin and J. S. Flier (2007). “Evidence for constitutive neural cell proliferation in the adult murine hypothalamus.” J Comp Neurol 505(2): 209–220.

44. Krabichler, Q., A. Lefevre, A. Kania, D. Hagiwara, K. Afordakos, K. Schönig, D. Bartsch and G. G (2023). Using a novel transgenic AVP-Cre rat to dissect arginine vasopressin circuits in the brain and their behavioral roles. Pan-American Physiological Sciences 2023, Puerto Varas, Chile.

45. Kumar, S. S. and P. S. Buckmaster (2007). “Neuron-specific nuclear antigen NeuN is not detectable in gerbil subtantia nigra pars reticulata.” Brain Res 1142: 54–60.

46. Lieber, C. S., S. Shaw and L. Van Waes (1978). “Alcoholism and alcoholic liver injury: new diagnostic and prognostic tests.” Arch Pathol Lab Med 102(8): 393–395.

47. Liu, R. Y., J. N. Zhou, W. J. Hoogendijk, J. van Heerikhuize, W. Kamphorst, U. A. Unmehopa, M. A. Hofman and D. F. Swaab (2000). “Decreased vasopressin gene expression in the biological clock of Alzheimer disease patients with and without depression.” J Neuropathol Exp Neurol 59(4): 314–322.

48. Lois, C. and A. Alvarez-Buylla (1994). “Long-distance neuronal migration in the adult mammalian brain.” Science 264(5162): 1145–1148.

49. Lois, C., J. M. Garcia-Verdugo and A. Alvarez-Buylla (1996). “Chain migration of neuronal precursors.” Science 271(5251): 978–981.

50. Lucas, C. H., M. Calvez, R. Babu and A. Brown (2014). “Altered subcellular localization of the NeuN/Rbfox3 RNA splicing factor in HIV-associated neurocognitive disorders (HAND).” Neurosci Lett 558: 97–102.

51. Markakis, E. A., T. D. Palmer, L. Randolph-Moore, P. Rakic and F. H. Gage (2004). “Novel neuronal phenotypes from neural progenitor cells.” J Neurosci 24(12): 2886–2897.

52. McPhail, L. T., C. B. McBride, J. McGraw, J. D. Steeves and W. Tetzlaff (2004). “Axotomy abolishes NeuN expression in facial but not rubrospinal neurons.” Exp Neurol 185(1): 182–190.

53. Mecawi, A. S. (2023). Multi-omics analyses of the hypothalamic- neurohypophysial system. Pan-American Physiological Sciences 2023, Ouerto Varas, Chile.

54. Migaud, M., M. Batailler, S. Segura, A. Duittoz, I. Franceschini and D. Pillon (2010). “Emerging new sites for adult neurogenesis in the mammalian brain: a comparative study between the hypothalamus and the classical neurogenic zones.” Eur J Neurosci 32(12): 2042–2052.

55. Ming, G. L. and H. Song (2011). “Adult neurogenesis in the mammalian brain: significant answers and significant questions.” Neuron 70(4): 687–702.

56. Pauza, A. G., A. S. Mecawi, A. Paterson, C. C. T. Hindmarch, M. Greenwood, D. Murphy and M. P. Greenwood (2021). “Osmoregulation of the transcriptome of the hypothalamic supraoptic nucleus: A resource for the community.” J Neuroendocrinol 33(8): e13007.

57. Ponti, G., P. Peretto and L. Bonfanti (2008). “Genesis of neuronal and glial progenitors in the cerebellar cortex of peripuberal and adult rabbits.” PLoS One 3(6): e2366.

58. Rankin, S. L., G. D. Partlow, R. D. McCurdy, E. D. Giles and K. R. Fisher (2003). “Postnatal neurogenesis in the vasopressin and oxytocin-containing nucleus of the pig hypothalamus.” Brain Res 971(2): 189–196.

59. Raymond, A. D., N. N. Kucherepa, K. R. Fisher, W. G. Halina and G. D. Partlow (2006). “Neurogenesis of oxytocin-containing neurons in the paraventricular nucleus (PVN) of the female pig in 3 reproductive states: puberty gilts, adult gilts and lactating sows.” Brain Res 1102(1): 44–51.

60. Sharif, A., C. P. Fitzsimons and P. J. Lucassen (2021). “Neurogenesis in the adult hypothalamus: A distinct form of structural plasticity involved in metabolic and circadian regulation, with potential relevance for human pathophysiology.” Handb Clin Neurol 179: 125–140.

61. Sladek, C. D. and W. E. Armstrong (1987). Effect of neurotransmitters and neuropeptides on vasopressin release. Vasopressin: principles & properties. D. M. Gash and G. J. Boer. New York, Plenum: 275–333.

62. Somogyi, P. and H. Takagi (1982). “A note on the use of picric acid-paraformaldehyde-glutaraldehyde fixative for correlated light and electron microscopic immunocytochemistry.” Neuroscience 7(7): 1779–1783.

63. Son, J. E., Z. Dou, K. H. Kim, S. Wanggou, V. S. B. Cha, R. Mo, X. Zhang, X. Chen, T. Ketela, X. Li, X. Huang and C. C. Hui (2021). “Irx3 and Irx5 in Ins2-Cre(+) cells regulate hypothalamic postnatal neurogenesis and leptin response.” Nat Metab 3(5): 701–713.

64. Swanson, L. W. (2004). Brain Maps: Structure of the Rat Brain. Oxford, Elservier Academic Press.

65. Tahara, A., J. Tsukada, Y. Tomura, T. Yatsu and M. Shibasaki (2011). “Vasopressin induces human mesangial cell growth via induction of vascular endothelial growth factor secretion.” Neuropeptides 45(2): 105–111.

66. Ugrumov, M. V. (2002). “Magnocellular vasopressin system in ontogenesis: development and regulation.” Microsc Res Tech 56(2): 164–171.

67. Unal-Cevik, I., M. Kilinc, Y. Gursoy-Ozdemir, G. Gurer and T. Dalkara (2004). “Loss of NeuN immunoreactivity after cerebral ischemia does not indicate neuronal cell loss: a cautionary note.” Brain Res 1015(1-2): 169–174.

68. van Eerdenburg, F. J., C. M. Lugard-Kok, S. J. Dieleman, M. M. Bevers and D. F. Swaab (1991). “Influence of gonadectomy and testosterone supplementation on the postnatal development of the vasopressin and oxytocin-containing nucleus of the pig hypothalamus.” Neuroendocrinology 54(6): 580–586.

69. van Eerdenburg, F. J., P. Poot, G. J. Molenaar, F. W. van Leeuwen and D. F. Swaab (1990). “A vasopressin and oxytocin containing nucleus in the pig hypothalamus that shows neuronal changes during puberty.” J Comp Neurol 301(1): 138–146.

70. van Eerdenburg, F. J. and D. F. Swaab (1991). “Increasing neuron numbers in the vasopressin and oxytocin containing nucleus of the adult female pig hypothalamus.” Neurosci Lett 132(1): 85–88.

71. van Eerdenburg, F. J. and D. F. Swaab (1994). “Postnatal development and sexual differentiation of pig hypothalamic nuclei.” Psychoneuroendocrinology 19(5-7): 471–484.

72. Wang, S., X. Li, M. Parra, E. Verdin, R. Bassel-Duby and E. N. Olson (2008). “Control of endothelial cell proliferation and migration by VEGF signaling to histone deacetylase 7.” Proc Natl Acad Sci U S A 105(22): 7738–7743.

73. Wigger, A., M. M. Sanchez, K. C. Mathys, K. Ebner, E. Frank, D. Liu, A. Kresse, I. D. Neumann, F. Holsboer, P. M. Plotsky and R. Landgraf (2004). “Alterations in central neuropeptide expression, release, and receptor binding in rats bred for high anxiety: critical role of vasopressin.” Neuropsychopharmacology 29(1): 1–14.

74. Wu, K. L., Y. Q. Li, A. Tabassum, W. Y. Lu, I. Aubert and C. S. Wong (2010). “Loss of neuronal protein expression in mouse hippocampus after irradiation.” J Neuropathol Exp Neurol 69(3): 272–280.

75. Yao, Y., A. B. Taub, J. LeSauter and R. Silver (2021). “Identification of the suprachiasmatic nucleus venous portal system in the mammalian brain.” Nat Commun 12(1): 5643.

76. Yoo, S. and S. Blackshaw (2018). “Regulation and function of neurogenesis in the adult mammalian hypothalamus.” Prog Neurobiol 170: 53–66.

77. Zhang, L. and L. E. Eiden (2019). “Two ancient neuropeptides, PACAP and AVP, modulate motivated behavior at synapses in the extrahypothalamic brain: a study in contrast.” Cell Tissue Res 375(1): 103–122.

78. Zhang, L., L. Guadarrama, A. A. Corona-Morales, A. Vega-Gonzalez, L. Rocha and A. Escobar (2006). “Rats subjected to extended L-tryptophan restriction during early postnatal stage exhibit anxious-depressive features and structural changes.” J Neuropathol Exp Neurol 65(6): 562-570.

79. Zhang, L. and V. S. Hernandez (2013). “Synaptic innervation to rat hippocampus by vasopressin-immuno- positive fibres from the hypothalamic supraoptic and paraventricular nuclei.” Neuroscience 228: 139–162.

80. Zhang, L., V. S. Hernandez, F. S. Estrada and R. Lujan (2014). “Hippocampal CA field neurogenesis after pilocarpine insult: The hippocampal fissure as a neurogenic niche.” J Chem Neuroanat 56: 45-57.

81. Zhang, L., V. S. Hernandez, B. Liu, M. P. Medina, A. T. Nava-Kopp, C. Irles and M. Morales (2012). “Hypothalamic vasopressin system regulation by maternal separation: its impact on anxiety in rats.” Neuroscience 215: 135–148.

82. Zhang, L., V. S. Hernández, D. Murphy, W. S. Young and L. E. Eiden (2021). Fine chemo-anatomy of hypothalamic magnocellular vasopressinergic system with an emphasis on ascending connections for behavioural adaptation. Neuroanatomy of Neuroendocrine Systems Á. D. Valery Grinevich, Springer- Nature.

83. Zhang, L., V. S. Hernandez, E. Vazquez-Juarez, F. K. Chay and R. A. Barrio (2016). “Thirst Is Associated with Suppression of Habenula Output and Active Stress Coping: Is there a Role for a Non-canonical Vasopressin-Glutamate Pathway?” Front Neural Circuits 10: 13.

84. Zhang, L., V. S. Hernandez, M. A. Zetter and L. E. Eiden (2020). “VGLUT-VGAT expression delineates functionally specialised populations of vasopressin-containing neurones including a glutamatergic perforant path-projecting cell group to the hippocampus in rat and mouse brain.” J Neuroendocrinol 32(4): e12831.

85. Zhao, C., W. Deng and F. H. Gage (2008). “Mechanisms and functional implications of adult neurogenesis.” Cell 132(4): 645–660.

86. Zhao, M., S. Momma, K. Delfani, M. Carlen, R. M. Cassidy, C. B. Johansson, H. Brismar, O. Shupliakov, J. Frisen and A. M. Janson (2003). “Evidence for neurogenesis in the adult mammalian substantia nigra.” Proc Natl Acad Sci U S A 100(13): 7925–7930.

87. Zhou, Y., Y. Su, S. Li, B. C. Kennedy, D. Y. Zhang, A. M. Bond, Y. Sun, F. Jacob, L. Lu, P. Hu, A. N. Viaene, I. Helbig, S. K. Kessler, T. Lucas, R. D. Salinas, X. Gu, H. I. Chen, H. Wu, J. E. Kleinman, T. M. Hyde, D. W. Nauen, D. R. Weinberger, G. L. Ming and H. Song (2022). “Molecular landscapes of human hippocampal immature neurons across lifespan.” Nature 607(7919): 527–533.

